# JASPER: fast, powerful, multitrait association testing in structured samples gives insight on pleiotropy in gene expression

**DOI:** 10.1101/2023.12.18.571948

**Authors:** Joelle Mbatchou, Mary Sara McPeek

**Author notes:** Corresponding author: Mary Sara McPeek.

## Abstract

Joint association analysis of multiple traits with multiple genetic variants can provide insight into genetic architecture and pleiotropy, improve trait prediction and increase power for detecting association. Furthermore, some traits are naturally high-dimensional, e.g., images, networks or longitudinally measured traits. Assessing significance for multitrait genetic association can be challenging, especially when the sample has population sub-structure and/or related individuals. Failure to adequately adjust for sample structure can lead to power loss and inflated type 1 error, and commonly used methods for assessing significance can work poorly with a large number of traits or be computationally slow. We developed JASPER, a fast, powerful, robust method for assessing significance of multitrait association with a set of genetic variants, in samples that have population sub-structure, admixture and/or relatedness. In simulations, JASPER has higher power, better type 1 error control, and faster computation than existing methods, with the power and speed advantage of JASPER increasing with the number of traits. JASPER is potentially applicable to a wide range of association testing applications, including for multiple disease traits, expression traits, image-derived traits and microbiome abundances. It allows for covariates, ascertainment and rare variants and is robust to phenotype model misspecification. We apply JASPER to analyze gene expression in the Framingham Heart Study, where, compared to alternative approaches, JASPER finds more significant associations, including several that indicate pleiotropic effects, some of which replicate previous results, while others have not previously been reported. Our results demonstrate the promise of JASPER for powerful multitrait analysis in structured samples.

## Introduction

In genome-wide association studies (GWAS), jointly analyzing multiple traits can give important insight about genetic architecture of complex traits [1–5] as well as improve predictions of comorbidity based on genetic information [6–8]. Multitrait analysis can also enhance the ability to detect genetic association even in the absence of pleiotropy [9–13]. In addition, some traits of interest for genetic analysis are naturally high-dimensional, such as those arising from medical imaging [14, 15] or longitudinally measured traits [16–18]. Furthermore, joint analysis of multiple traits with multiple rare (or common) variants can improve the ability to detect gene-based associations [3, 14].

When testing for association between a multi-dimensional trait (or a set of traits) and multiple genetic variants, obtaining an accurate assessment of significance can be challenging, especially when there are related individuals or population sub-structure in the sample. Failing to adequately adjust for sample structure when it is present can lead to inflated type 1 error rates as well as power loss. Typical ways to assess significance for an association test statistic include asymptotic approximations [10, 12, 19, 20] and permutation tests [21–26]. Asymptotic approximations can be fast to compute, but can break down with high-dimensional outcomes [27, 28] or with small sample sizes [19, 29] and typically rely on correct phenotype model specification. These problems can be avoided by using retrospective association testing [16, 30–34] with asymptotic assessment of significance. However, when used with multiple genetic variants [32] retrospective association tests that use asymptotic assessments of significance can be overly sensitive to estimation of the covariance matrix for the genetic variants, which may not be accurately estimated, particularly with rare variants. Permutation tests can be used to solve the above problems. While most permutation approaches require no population or pedigree structure in the sample, exceptions do exist [21, 22, 24, 26]. However, the main drawback of permutation tests is that they are computationally very expensive. A third approach [28, 35, 36] uses a moment-matching procedure to approximate the empirical distribution of all possible permutations of the test statistic under the null hypothesis of no association. This approximation to a permutation test has the robustness of a permutation test, but with fast computational speed, because analytical expressions of the first three moments needed for moment-matching have been previously derived [37]. However, currently this approach is only valid when either (1) there is no population or pedigree structure among the individuals or (2) the structure in the data is so simple that it can be perfectly captured by a set of ancestry informative vectors that are included as covariates in the model, which is not realistic for many samples that include more complex population or pedigree structure or cryptic relatedness.

We consider the problem of testing for association between a set of traits and a set of genetic variants in the presence of population and/or pedigree structure. We propose a very general approach to assessing significance in structured samples: JASPER (Joint Association analysis in Structured samples based on approximating a PERmutation distribution), which extends the fast moment-matching approximation method to account for sample structure. Underlying JASPER is a new analytical approximation to the first three moments of the permutation distribution that is general enough to allow for sample structure. JASPER is applicable to essentially arbitrary phenotype categories (e.g., quantitative, case-control, count, high-dimensional) and is designed to be robust to phenotype model misspecification. It allows for covariates and uses a genetic relatedness matrix (GRM) to correct for population structure. JASPER is applicable to to a wide range of association test statistics, including methods for associating a set of rare variants with a single continuous trait (SKAT [19], famSKAT [38], generalized SKAT [29] and MONSTER fixed-*ρ* [39]) or with a single binary trait [32], as well as kernel-based multitrait methods for testing association with a set of rare or common variants, when these are used with a linear or weighted linear kernel for the genotypes, such as DKAT [28], MSKAT [10], GAMuT [20], the kernel-based test statistics that are combined into the omnibus test in Multi-SKAT [12], wSKAT [40], and a kernel-based method for genetic analysis of the microbiome [41]. Simple special cases also include single-trait, single variant association tests for binary traits [33, 34] and for quantitative traits (score and Wald tests for association in a linear mixed model).

JASPER is potentially applicable to a wide range of association testing problems. We illustrate how to apply JASPER to (1) start with a multitrait, multi-variant association test that does not adequately account for population structure, and “robustify” it, so that it properly accounts for population structure; (2) derive a fast, robust assessment of significance for a multitrait, multi-variant association test that allows for population structure, in a high dimensional setting in which an asymptotic assessment of significance breaks down and (3) start with a rare-variant binary trait association test that does not adequately account for population structure, and “robustify” it, so that it properly accounts for population structure. We demonstrate through simulation studies in structured samples that JASPER improves both type 1 error and power compared to both permutation methods and asymptotic assessments of significance across a wide range of sample structure settings and trait models, with particularly notable improvement in power when the number of traits is large. We also show that JASPER is both computationally fast and robust against phenotype model misspecification. We apply JASPER to analyze genetic regulation of gene expression levels within biological pathways in data from the Framingham Heart Study (FHS), which involves multitrait mapping of hundreds of traits simultaneously with multiple genetic variants in a sample with pedigree structure.

## Material and Methods

We consider a relatively general context in which *k* traits have been measured on *n* individuals, and association is being tested jointly between the *k* traits and a set of *m* genetic variants. For example, the *k* traits might be gene expression values for *k* genes in a biological pathway, and the set of *m* genetic variants being tested might consist of variants within a single gene. In this work, we explicitly consider both quantitative and binary traits, but the type of trait that can be accommodated is completely general, and the genetic variants could be common or rare. Population substructure, admixture and relatedness, either known or cryptic, can be present in the sample. Covariates, if available, can also be accommodated in the analysis.

We define a very general class of association test statistics, which we call the JASPER class, that includes many popular single- and multitrait test statistics for testing association with sets of rare or common variants. We show how to use the JASPER method to develop a fast, robust assessment of significance for any test statistic in the JASPER class. This allows us to increase power and improve control of false positives in genetic association analysis across a wide range of contexts. For example, with JASPER, we are able to take into account population and pedigree structure to quickly and correctly assess significance for a test statistic even if (1) the test statistic itself does not properly account for population structure or is based on a misspecified phenotype model or (2) the test statistic is based on a large number of traits (and/or variants) for which asymptotic approximations break down or (3) the sample size is small, in which case the asymptotic approximations break down.

### Basic notation

Let *Y* denote the *n* ×*k* matrix of *k* traits among *n* subjects, *G* denote the *n* ×*m* genotype matrix for the *m* genetic variants being simultaneously tested (out of a potentially much larger set of genomewide genetic variants on which data are available), with *G*_*ij*_ being the genotype (e.g., minor allele count: 0, 1, or 2), for the *i*th individual at the *j*th variant. Let *X* be an *n* ×*c* matrix of covariates, including an intercept. Let *X*_*G*_ be an 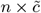 sub-matrix of *X* consisting of the confounding covariates, e.g., ancestry informative vectors such as principal components (PCs) and any other covariates that are associated with *G* as well as with *Y*, where 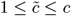 and *X*_*G*_ consists of a subset of the columns of *X*, including, at a minimum, the intercept column, and at a maximum, *X*_*G*_ = *X*. We assume that any confounding covariates that are available and included in *X* are also included in *X*_*G*_. We define 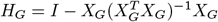, which is the projection that maps a vector to its residuals after regressing out *X*_*G*_.

### A general class of genetic association test statistics

We consider a general class of genetic association test statistics that can be written in the form

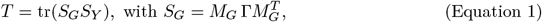

where tr(·) denotes the trace operator of a matrix, *S*_*Y*_ is a symmetric, positive semi-definite *n* ×*n* matrix (the “phenotype kernel”), that is a function of *Y* and *X, M*_*G*_ is an *n* ×*m* matrix, and Γ is an *m* ×*m* symmetric, positive semi-definite matrix, so that *S*_*G*_ is also a symmetric, positive semi-definite *n* ×*n* matrix (the “genotype kernel”). In the context of genetic association testing, we specifically assume that *M*_*G*_ = *MG*, where *M* is an *n* ×*n* matrix that can be a function of *X* and is assumed to be non-random given *X*. We further assume that *MX*_*G*_ = 0. In the example test statistics described in detail in this paper, *M* can be taken to be *M* = *H*_*G*_. We assume that under the null hypothesis, *S*_*G*_ and *S*_*Y*_ are conditionally independent given *X*. Many widely-used genetic association test statistics are in this class, as detailed in the **Introduction**.

The genetic association test statistics in Equation 1 can be considered as a special case of a more general JASPER class of statistics to test association between two sets of generic variables, *G* and *Y*, not necessarily genotype and phenotype, where *M*_*G*_ is allowed to be more general than just a linear function of *G*, and where the conditions of Proposition 1 in **Appendix A** are required to hold. These conditions basically ensure that we have some way to decorrelate the rows and columns of *S*_*G*_, which is required to be able to use the JASPER method to assess significance in the presence of dependence between the samples (where, in the genetic association context, the dependence we need to correct for corresponds to population and/or pedigree structure). Interestingly, for JASPER, it is not necessary to know or estimate the correlation among traits or among genetic variants, in the sense that these values are not needed to ensure correct type 1 error of the test (though one can choose to incorporate this additional information into the test statistic for the purpose of increasing power).

The JASPER class is related to the RV coefficient class of test statistics (e.g., [42]), where association is tested between multidimensional variables *A* and *B* using a test statistic of the form

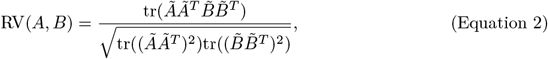

where *Ã* = *HA*, 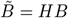, *H* = *I* − *n*^−1^*J*_*n*_, and *J*_*n*_ is an *n* ×*n* matrix with every element equal to 1. If we define *G* to be *A* and *Y* to be *B*, and set 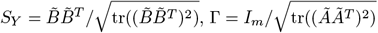, where *I*_*m*_ is the identity matrix of dimension *m*, and *M*_*G*_ = *Ã*, then we can write RV(*A, B*) in the form of Equation 1. However, the JASPER class is more general than the RV class in a very useful way: the JASPER class allows for population and pedigree structure among the individuals, whereas the RV class does not. Similarly, JASPER is closely related to the DKAT statistics [28], which are of the form in Equation 2 where *AA*^*T*^ is taken to be any positive semi-definite genotype kernel and *BB*^*T*^ is taken to be any positive semi-definite phenotype kernel. The DKAT methods require independence of individuals, and JASPER can be considered an extension of DKAT to the case when there is population and/or pedigree structure, where we put some additional requirements on one of the two kernels (in this case, *S*_*G*_), to ensure that we have a way to decorrelate its rows and columns. In all the worked examples in this paper, we are using a weighted linear genotype kernel of the form *S*_*G*_ = *H*_*G*_*G*Γ*G*^*T*^ *H*_*G*_, but any phenotype kernel can be used for *S*_*Y*_.

### A very basic example

The association tests we focus on in this work involve joint association testing with either multiple traits or multiple genetic variants, or both. However, to start, it can be helpful to understand the JASPER class by considering a simpler example. A fairly simple example of a test statistic of this type in the special case *k* = *m* = 1 would be either the Wald test statistic or score test statistic for association between a genetic variant and a phenotype in a linear mixed model (LMM) of the form

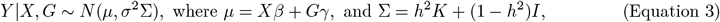

where *Y* is *n* ×1 (i.e., *k* = 1), *X* is *n* ×*c*, and *G* is *n* ×1 (i.e., *m* = 1), all defined as before, *β* is a *c* ×1 vector of unknown coefficients, *K* is an *n×n* GRM or kinship matrix, *h*^2^ ∈ [0, 1) and *σ*^2^ > 0 are unknown parameters, and *γ* is the unknown scalar parameter of interest. To test the null hypothesis *H*_0_ : *γ* = 0, both the squared Wald and score test statistics would be of the form in Equation 1, where *M*_*G*_ = *MG*, with *M* = *H*_*G*_, 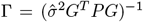 and *S*_*Y*_ = *PY Y* ^*T*^ *P* with 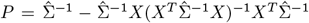, where *ĥ*^2^ and 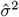are estimators of *h*^2^ and *σ*^2^, and 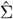 is Σ with *ĥ*^2^ plugged in. The Wald and score test statistics differ only in the choice of estimators *ĥ*^2^ and 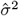. For our purposes, we could ignore the scalar multiple 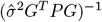 (which will be unchanged by permutations of the rows and columns of *S*_*G*_, or of its transformed version 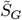, defined later, and so will have no impact), and instead take Γ = 1. Note that instead of *M* = *H*_*G*_, we could take 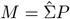, because the identities 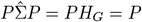 ensure that these two choices of *M* will lead to the same value of *T*. An even simpler example would be the t-test for significance of *γ* in a simple linear regression of *Y* on *X* and *G*, for which *M*, Γ and *S*_*Y*_ could be defined as above with 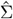 replaced by the identity matrix.

### A fast, robust assessment of significance when there is no sample structure

In the absence of population structure or related individuals in the sample, one could use a permutation test to assess significance of a test statistic in the JASPER class as follows: Randomly permute the *n* rows and columns of the matrix *S*_*G*_ (where we apply the same permutation to both rows and columns) *p* times to obtain permutation replicates 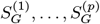 and calculate 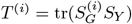 for the *i*th permutation replicate to generate an empirical null distribution for *T* based on *T* ^(1)^, …, *T* ^(*p*)^. Note that applying a permutation *σ* to both the rows and columns of *S*_*G*_ gives the same result as (1) applying the permutation *σ*^−1^ to both the rows and columns of *S*_*Y*_ or (2) applying the permutation *σ* to the rows of *M*_*G*_. From, we can see that the permutation test based on permuting rows and columns of *S*_*G*_ and that based on permuting rows and columns of *S*_*Y*_ are equivalent. From (2), we can see that the proposed permutation test would be appropriate if, under the null hypothesis of no association, the distribution of *T* were assumed to be invariant to permutation of the rows of *M*_*G*_. A sufficient condition for this would be that, under the null hypothesis of no association, *S*_*G*_ and *S*_*Y*_ are independent (or conditionally independent given some covariates *X*) and the rows of *M*_*G*_ are exchangeable (meaning that the distribution of *M*_*G*_ under the null hypothesis is unchanged by permutation of the rows).

As a faster alternative to performing actual permutations, one could instead approximate the permutation distribution of *T* conditional on *S*_*Y*_ and *S*_*G*_ by a Pearson type III distribution using moment-matching based on exact analytical calculations of the first three moments of *T* [28, 35–37, 42]. Previous work [42] has demonstrated the effectiveness of the Pearson type III distribution to adequately capture the skewness component of permutation distributions. A key advantage of this approach is that it eliminates the need to explicitly carry out the permutations themselves and thus drastically decreases the computational cost involved in assessing significance. However, this approach is currently valid only when there is no sample structure. In the next 4 subsections, we describe our extension of this approach to allow population and pedigree structure in the sample.

### JASPER: A fast robust assessment of significance when sample structure is present

We instead consider a more general situation in which there is sample structure, e.g., population structure and/or relatedness, so that under the null hypothesis, the rows of *M*_*G*_ are not exchangeable. In Proposition 1 in **Appendix A**, we define transformed kernels 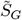 and 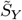 and show that for test statistics *T* in the JASPER class, we can rewrite *T* as

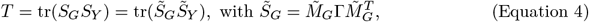

where, under the null hypothesis, 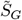 and 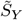 are conditionally independent given covariates *X*, and the rows of 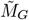 are second order exchangeable under the null hypothesis, so that it would be justified to obtain an empirical null distribution for *T* by permuting the rows and columns of 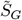, using the same permutation for the rows and columns (which is equivalent to permuting the rows of the matrix 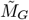), *p* times and recalculating *T* for each. Construction of 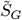 and 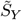 is described in sub-section **Decorrelation of** *S*_*G*_ **in JASPER** below. Even better, in sub-section **Extension of fast approximation of p-values to allow for population structure**, we also show how to avoid actually doing any permutations by instead calculating the approximate null permutation distribution of *T*. This requires a new analytical approximation to the first three moments of the permutation distribution that is general enough to allow for sample structure, which is given in Proposition 2 in **Appendix B**, leading to a fast p-value calculation.

### Assumptions on null conditional mean and variance of genotype matrix

In later sub-sections, we show how to apply JASPER to (1) start with a multitrait, multi-marker association test that does not adequately account for population structure, and “robustify” it, so that it properly accounts for population structure; (2) derive a fast, robust assessment of significance for a multitrait, multi-marker association test that allows for population structure, in a high dimensional setting in which an asymptotic assessment of significance breaks down and (3) start with a rare variant binary trait association test that does not adequately account for population structure, and “robustify” it, so that it properly accounts for population structure. In all three of these applications of JASPER, we make certain basic assumptions on the first and second moments of the genotype matrix, which are that under the null hypothesis of no association,

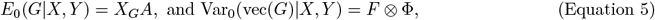

where *A* is a 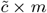 matrix of unknown coefficients, vec(·) is the vectorization operator that turns a matrix into a vector by stacking its columns, *F* is an *m* ×*m* positive semi-definite matrix that represents the linkage disequilibrium structure (i.e., covariance) among the *m* genetic variants and Φ is an *n* ×*n* positive semi-definite matrix that represents the genetic covariance among the *n* individuals. We typically estimate Φ by *K*, where *K* is either the GRM based on genomewide variants or the kinship matrix. In the special case when *m* = 1 and *X*_*G*_ = *X*, these moment conditions have previously been used in retrospective association tests with single variants [16, 31, 33, 34].

An exciting aspect of our method is the lack of assumptions required for it. Specifically, (1) we do not need to make any assumptions about the distribution of *Y*, except that we assume that under the null hypothesis, *Y* is conditionally independent of *G* given *X*, and (2) we do not need to make any assumptions about the form of *F*, the covariance matrix among the tested SNPs, which we do not need to estimate at all. Assumptions about the distribution of *Y* are problematic because in practice, important features of the true phenotypic distribution are generally unknown. Estimation of *F* can be particularly problematic when some of the *m* variants are rare, because there is often not sufficient information available, either within the data set or from a suitable reference panel, to obtain accurate estimation of *F*. In contrast, our method works well as long as a suitable estimate of Φ is available.

The JASPER method does not depend specifically on the assumptions in Equation 5, though some first and second moment assumptions are required in order to allow us to decorrelate one of the two kernels, as described in Proposition 1. The assumptions in Equation 5 could be modified as desired, as long as the conditions of Proposition 1 are met and 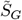and 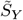 are defined accordingly.

### Decorrelation of *S*_*G*_ in JASPER

Using the assumptions of the previous subsection, for a genetic association test in which *M*_*G*_ = *MG*, we can apply Proposition 1 (**Appendix A**) to decorrelate the rows and columns of *S*_*G*_, with corresponding changes in *S*_*Y*_, to obtain the new matrices 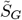 and 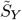 of Equation 4. We now describe how this is done. Recall that we have assumed *MX*_*G*_ = 0, so it follows from this and Equation 5, using standard identities for the vec operator applied to a matrix product, that

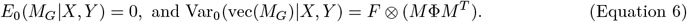

Then the needed matrix *V*_*r*_ in Proposition 1 is equal to *M* Φ*M* ^*T*^, the estimate of *V*_*r*_ is taken to be 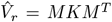, where *K* is the GRM or kinship matrix, and the conditions of Proposition 1 (using Remark 6) are trivially seen to be satisfied. Then to construct 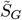 and 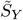, we first let 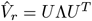 be the eigendecomposition of the *n* ×*n* matrix 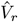. Let *ñ* be the rank of 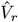, which would typically be 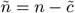, and define Λ_1*/*2_ to be the *n*×*ñ* matrix consisting of the *ñ* nonzero columns of the *n*×*n* matrix Λ^1*/*2^, and define Λ_−1*/*2_ to be the *n* ×*ñ* matrix consisting of the *ñ* non-zero columns of the *n* ×*n* matrix (Λ^−^)^1*/*2^, where Λ^−^ is the Moore-Penrose generalized inverse of Λ. (Λ_−1*/*2_ could be constructed from Λ_1*/*2_ by replacing each of the *ñ* non-zero elements, *e*_*i*_, by 1*/e*_*i*_ and leaving the zero elements unchanged.) Then we define 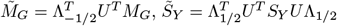, and 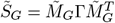. Note that 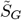 and 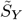are of dimension *ñ < n*.

By Proposition 1, we have now successfully decorrelated the rows and columns of *S*_*G*_ to form 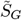 and a corresponding 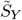 in such a way that Equation 4 holds, and we can now apply an actual or approximate permutation test to assess significance of *T*. In the next subsection we describe our fast approximation to a permutation distribution for *T*.

### Extension of fast approximation of p-values to allow for population structure

In principle, we could assess the significance of *T* by computing its null distribution on all possible permutations of the rows of 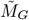, or on a random sample of permutations. As this approach is computationally intensive, we propose to instead use a moment-matching procedure based on approximating this distribution with a Pearson type III distribution using exact analytical calculations of the first three moments of *T*. Such an approach has previously been used only in the case when all sampled individuals are assumed to be independent, and in Proposition 2 (**Appendix B**) we prove a more general result that is needed to extend the approximation to allow for sample structure. 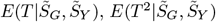 and 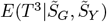 are derived in Proposition 2 of **Appendix B** and **Supplemental Methods**, and from these, the mean, variance and skewness of *T* are computed as

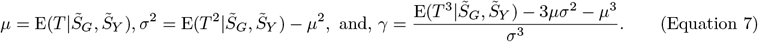

Using these, we approximate the null permutation distribution of *T* by a Pearson type III distribution with scale parameter *a* = *γσ/*2,, shape *b* = 4*/γ*^2^, and location parameter 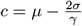, where the probability density function is

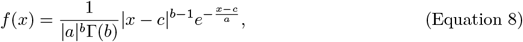

for *a*≠0, *b* > 0 and 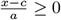. We then obtain a p-value analytically by calculating 1 − *F* (*T*), where *F* is the cdf corresponding to density *f*, and *T* is the observed value of the test statistic.

### Use of JASPER to “Robustify” a Multitrait, Multi-Marker Association Test

We consider the problem of testing association jointly between a set of multiple quantitative traits and a set of multiple variants. We start with a simple multitrait, multi-marker, association test statistic, *T*, that either (1) ignores population structure or (2) only partially accounts for it by including some PCs as covariates. Using the JASPER method for assessing significance, we show how to “robustify” *T*, so that the resulting test based on *T* is correctly calibrated even when both population structure and pedigree structure are present.

The statistic *T* is constructed based on a LMM with the following form:

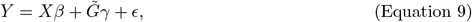

where *Y* is *n* ×*k* and *X* is *n* ×*c*, both defined as before; *β* is a *c* ×*k* matrix of fixed effects; vec(*ϵ*) ∼ *N* (0, *V*_*e*_ ⊗*I*_*n*_), where *V*_*e*_ is an unknown *k×k* trait covariance matrix and *I*_*n*_ is the *n×n* identity matrix; 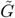 is a *n* ×*m* genotype matrix that has been centered by regressing out the confounding covariates *X*_*G*_ and standardized so that each column has mean 0 and variance 1; *γ* is an *m* ×*k* matrix of random effects for the tested variants, where we do not specify the full distribution for *γ*, but only require that *E*[vec(*γ*)] = 0 and Var[vec(*γ*)] = *τ* ^2^*V*_*g*_ ⊗ *W*, where *V*_*g*_ is a *k* ×*k* covariance matrix that is either pre-specified or is set equal to *V*_*e*_, *W* is a pre-specified *m* ×*m* positive definite “weight matrix”, and *τ* ^2^ is an unknown scalar. Then for the model in Equation 9, we have 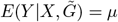 and 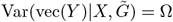, where *µ* = *Xβ* and

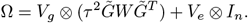

We test the null hypothesis *H*_0_ : *τ* ^2^ = 0, which corresponds to a joint test of association between the set of *k* traits and the set of *m* SNPs. The score test statistic, *T*, for this null hypothesis can be written in the form of Equation 1, i.e., *T* = tr(*S*_*G*_*S*_*Y*_), where 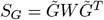, and

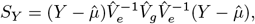

where 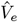 is the estimate of *V*_*e*_ under the null hypothesis; 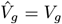 if *V*_*g*_ is pre-specified and 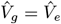 if *V*_*g*_ = *V*_*e*_ is assumed; and 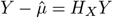, with *H*_*X*_ = *I* − *X*(*X*^*T*^ *X*)^−1^*X*^*T*^. Assuming that 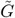was formed as 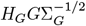, where 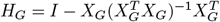 is the *n* ×*n* matrix used to center *G* and regress out any confounding covariates and Σ^−1*/*2^ is the *m* ×*m* matrix used to standardize *G*, where Σ_*G*_ is diagonal with *j*th diagonal element equal to the estimated conditional variance of *G*_*j*_ given *X*_*G*_, then we can write

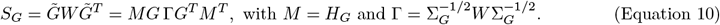

Note that in writing down the score test statistic for the variance component score test, we have used the common convention of discarding a term that subtracts off the null mean of the test statistic, because this term would have no impact on the JASPER assessment of significance.

With the definition of *M* in Equation 10, and assuming we have calculated a GRM or kinship matrix *K* to use as an estimate of Φ, we can follow the recipe in the subsection **Decorrelation of** *S*_*G*_ **in JASPER**, which involves taking an eigendecomposition of *MKM* ^*T*^ and using it to obtain 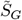 and 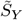. From 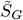 and 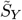, the JASPER procedure can directly obtain a p-value for *T* by using Proposition 2 to obtain the needed moments for the Pearson type III approximation to the distribution of *T*.

### Useful special cases I

In the special case of Equation 9 when we assume *V*_*g*_ = *V*_*e*_, we get

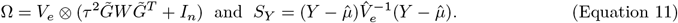

We define *T*_1_ to be the test statistic calculated from the *S*_*G*_ in Equation 10 and the *S*_*Y*_ in Equation 11, where *W* = Σ_*G*_ so Γ = *I*_*m*_, and

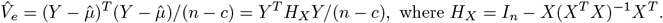

In **Results**, we use simulations to assess the ability of JASPER to robustify *T*_1_ in the presence of population and pedigree structure.

In the special case of Equation 9 when *k* = 1, without loss of generality, we can take *V*_*g*_ = 1, *V*_*e*_ reduces to the scalar *σ*^2^, *S*_*G*_ is the same as in Equation 10, and we get 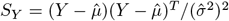, where the scalar multiple 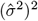 can be neglected in JASPER because it is unaffected by permutations of rows and columns of 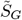. The SKAT statistic [19] and generalized or unified SKAT statistic [29] are both of this form and differ only in the choice of the matrix *W*.

### Use of JASPER for High-Dimensional Testing When Asymptotics Break Down

We again consider the problem of testing association jointly between a set of multiple quantitative traits and a set of multiple variants. Even when population structure is appropriately incorporated into the test statistic, and the phenotype is correctly modeled, asymptotic assessments of significance still tend to break down when the number of traits, *k*, grows large. We show how to use the JASPER method for assessing significance to obtain correctly calibrated association test statistics in structured samples even when the number of traits is large.

We consider a LMM with the following form:

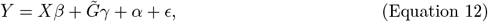

where *Y*, *X*, 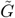, *β*, and *ϵ* are the same as in Equation 9; *α* represents additive polygenic random effects with vec(*α*) ∼ *N* (0, *V*_*a*_ ⊗ *K*), where *V*_*a*_ is an unknown *k* ×*k* positive definite matrix representing trait covariance due to additive polygenic effects and *K* is an *n* ×*n* GRM or kinship matrix; *γ* is an *m* ×*k* matrix of random effects for the tested variants, where we do not specify the full distribution for *γ*, but only require that *E*[vec(*γ*)] = 0 and Var[vec(*γ*)] = *τ* ^2^*V*_*g*_ ⊗*W*, where *V*_*g*_ is a *k×k* covariance matrix that is either pre-specified or is set equal to either *V*_*a*_ or *V*_*e*_, *W* is a pre-specified *m* ×*m* positive definite “weight matrix”, and *τ* ^2^ is an unknown scalar. Then for the model in Equation 12, we have 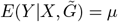 and 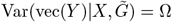, where *µ* = *Xβ* and

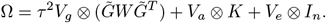

We test the null hypothesis *H*_0_ : *τ* ^2^ = 0, which corresponds to a joint test of association between the set of *k* traits and the set of *m* SNPs. The score test statistic for this null hypothesis can be written in the form of Equation 1 (see **Supplemental Methods** for proof), where *S*_*G*_ is the same as in Equation 10, and 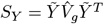, where 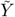 is the *n* ×*k* matrix such that

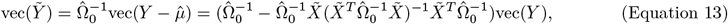

where 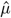 is the *n* ×*k* matrix whose vec is given by 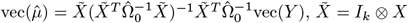 and 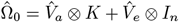, where 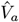 and 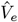 are the estimates of *V*_*a*_ and *V*_*e*_ under the null hypothesis. If *V*_*g*_ was pre-specified, then 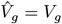. Otherwise, if *V*_*g*_ was set equal to *V*_*a*_ or *V*_*e*_, then 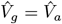 or 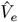, respectively.

Thus, we can define 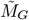 and 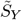 as in Proposition 1 and rewrite our test statistic as given in Equation 4, where under the null hypothesis, 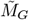 and 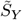are conditionally independent given *X*, and the rows of 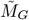are second-order exchangeable under the null hypothesis. Then we can apply JASPER’s fast approximation to the permutation distribution given in subsection **Extension of fast approximation of p-values to allow for population structure**.

### Useful special cases II

In our simulations, we consider the special case of Equation 12 in which *V*_*g*_ is set equal to *V*_*e*_, and *W* = Σ_*G*_, and we define test statistic *T*_2_ by *T*_2_ = tr(*S*_*G*_*S*_*Y*_), where 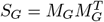, with *M*_*G*_ = *H*_*G*_*G*, and 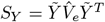. We use *T*_2_ in simulation studies to evaluate the ability of JASPER to correctly assess significance for high-dimensional testing when asymptotics break down.

In the special case of Equation 12 when *k* = 1, *S*_*G*_ is the same as in Equation 10, and without loss of generality, we can take *V*_*g*_ = 1, in which case we get 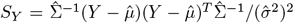, where Ω = *σ*^2^Σ with Σ defined in Equation 3, and where 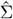 is Σ with *ĥ*^2^ plugged in. The resulting *S*_*Y*_ is the same as the one for the LMM with *k* = 1 and *m* = 1 in the subsection **A very basic example**, except for the scalar multiple 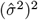, which, like all scalar multiples applied to the test statistic *T*, can be neglected in JASPER because it is unaffected by permutations of rows and columns of 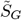. The famSKAT and MONSTER fixed-*ρ* statistics are both of this form and differ only in the choice of the symmetric matrix *W*.

### Use of JASPER to “Robustify” a binary-trait, rare-variant association test

The JASPER method is widely applicable. In this subsection, we consider the case of rare-variant association testing of a binary trait when population and pedigree structure are present in the sample. Dealing with multiple rare variants and population structure in the context of binary traits can be methodologically challenging. First, direct application of LMM approaches to binary traits suffers from power loss when covariate effects are strong [33, 34]. Generalized linear mixed-model methods [43] could, in principle, provide a solution to this problem, but in practice the penalized quasi-likelihood estimation methods that make them computationally feasible are too inaccurate to provide reliable type 1 error control [44]. Furthermore, test statistics based on binary trait models can be sensitive to model misspecification and ascertainment when a prospective assessment of significance is used [33, 43]. We start with a simple, binary-trait, rare-variant, association test statistic *T*_3_ that either (1) ignores population structure or (2) only partially accounts for it by including some PCs as covariates. Using the JASPER method for assessing significance, we show how to “robustify” *T*_3_, so that the resulting test based on *T*_3_ is correctly calibrated even when both population structure and pedigree structure are present.

For a single binary trait, we use the test statistic *T*_3_ which is based on a logistic mixed model [29, 45] in which

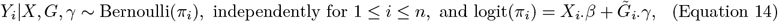

where *Y* is *n* ×1, *X* is *n* ×*c* and 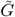 is *n* ×*m*, all defined as before; *X*_*i·*_ is the *i*th row of *X, β* is as defined before, 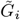*·* is the *i*th row of 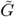, *γ* is a vector of length *m* of random effects for the tested variants, where we do not specify the full distribution for *γ*, but only require that *E*(*γ*) = 0 and Var(*γ*) = *Wτ* ^2^, where *W* is a pre-specified *m* ×*m* positive definite weight matrix and *τ* ^2^ is an unknown scalar. *T*_3_ corresponds to a score test for *H*_0_ : *τ* ^2^ = 0 and can be written in the form of Equation 1, where *S*_*G*_ has the same form as in Equation 10 and 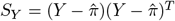, where 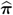 is the estimate of ***π*** = (*π*_1_, …, *π*_*n*_)^*T*^ obtained by fitting the logistic model logit(*π*_*i*_) = *X*_*i·*_*β*, which is what Equation 14 reduces to when *H*_0_ is true.

### Simulation studies

The JASPER method for assessing significance is potentially applicable to a wide range of contexts for association testing. To evaluate JASPER in simulations, we consider two different types of association testing: (1) multivariate association testing between a set of quantitative traits and a set of SNPs, and association testing between a single binary trait and a set of SNPs (either rare or common variants). Population and pedigree structure are simulated among the sampled individuals, and covariates are included. We consider 3 different test statistics, and for each, we compare JASPER to other methods of assessing significance. We evaluate the ability of JASPER to correctly control type 1 error, even when the statistic to which it is applied either ignores the population and pedigree structure or only partially accounts for it, in which case other methods to assess significance are expected to fail. We also evaluate the ability of JASPER to provide correct type 1 error control when assessing significance with high-dimensional traits, when asymptotic assessments of significance can break down, even if population and pedigree structure is appropriately accounted for. In addition, we compare the power of JASPER to that of other methods in these settings. In all settings, there is model misspecification in the fitted model, so that we can better assess how JASPER and the other methods would perform in realistic situations. Genotypes, traits and covariates are simulated under multiple sample structure settings and trait models, where we simulate *k* = 1, 2, 5, 50 or 100 traits.

#### Simulation settings: Population and pedigree structure and genotypes

The settings of population structure we consider are: (1) no population structure, (2) 2 sub-populations, and (3) 2 sub-populations with admixture, and each of these can be combined with pedigree structure or not. We also sample either rare or common variants. When pedigree structure is simulated, 46 three-generation pedigrees, each with 22 individuals are simulated, for a total of 1,012 sampled individuals. Genotypes or haplotypes are drawn for each founder, and gene-dropping is used to simulate the genotypes or haplotypes for the non-founder individuals in each pedigree. In the settings with no pedigree structure, 1,000 individuals are sampled at random from the structured population.

For common variants with 2 sub-populations, we use the 2-sub-population Balding-Nichols model [46] with *F* = .01 and ancestral allele frequencies for SNPs drawn independently and uniformly on (.2, .8). For common variants with 2 sub-populations with admixture and unrelated individuals, the allele frequency for an individual *i* is given by *a*_*i*_*p*_1_ + (1 − *a*_*i*_)*p*_2_, where *p*_1_ and *p*_2_ are the sub-population allele frequencies obtained from the Balding-Nichols model and *a*_*i*_ is the admixture proportion for individual *i*, where *a*_*i*_ is drawn from a Uniform(0, 1) distribution, i.i.d. across individuals. When pedigree structure is combined with setting (2) of population structure, 23 pedigrees are assigned to sub-population 1, and 23 are assigned to sub-population 2, and the founders of each pedigree are assumed to be sampled at random from the corresponding sub-population. When there is pedigree structure with no population structure, the founders of each pedigree are assumed to be drawn at random from the population. When pedigree structure is combined with setting (3) of population structure, an admixture proportion is sampled independently for each of the 46 pedigrees, and the founders are each assumed to have the corresponding admixture proportion. For rare variants, we simulate *m* = 50 rare variants with a common allele frequency *p*, where *p* = .1, .01, .005, or .001, where the rare variants are assumed to be correlated with a simple compound symmetric correlation structure [32], where the common correlation *ρ*_*g*_ takes value 0, .5 or .9. An exception is that for the case when *p* = .001, we did not use *ρ*_*g*_ = .9, because this setting is insufficiently polymorphic (as it results in a single haplotype with frequency > .99). For each simulation setting, in addition to the *m* tested variants, a kinship matrix *K* is simulated based on 10^5^ common variants.

#### Covariate and Trait Models

Three observed covariates are simulated: sex, age and a continuous covariate with the standard Gaussian distribution. (See **Supplemental Methods** for further details.) Additional covariates are used to generate the phenotypes, but are assumed to be unobserved in the analysis. These additional, unobserved covariates are (1) genotypes of two major causal variants, *M*_1_ and *M*_2_, where each of these is a vector of length *n*, and where these are assumed to be common variants simulated as described in the previous sub-section and (2) *a*, which is a vector of length *n* whose *i*th element is the proportion of ancestry from sub-population 1 for individual *i*, where this covariate only exists in the 2-sub-population and 2-sub-population with admixture settings.

We consider two models to generate the traits given the genotype and covariate data, where Model I is for simulating a set of *k* quantitative traits and Model II is for simulating a binary trait. In Model I, we generate the set of *k* quantitative traits as

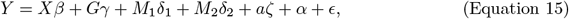

where *X* is *n* ×4 and consists of an intercept column and columns corresponding to the three observed covariates described above; *Y*, *G, M*_1_, *M*_2_, *a, β, γ, α*, and *ϵ* are as defined before, and *δ*_1_, *δ*_2_, and *ζ* are 1 ×*k* vectors of coefficients for the fixed effects of the unobserved covariates, *M*_1_, *M*_2_ and *a*, respectively. Recall that vec(*α*) ∼ *N* (0, *V*_*a*_ ⊗ *K*) and vec(*ϵ*) ∼ *N* (0, *V*_*e*_ ⊗ *I*_*n*_), where *V*_*a*_ and *V*_*e*_ are *k* ×*k* covariance matrices. We consider two different models for *V*_*a*_ and *V*_*e*_, depending on how many traits are being simulated. In the case when *k* ≤ 10, we use a compound symmetric model in which 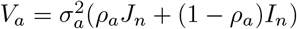 and 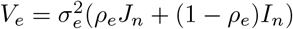, where *ρ*_*a*_ represents the genetic correlation between traits and *ρ*_*e*_ represents the residual correlation between traits. We fix *ρ*_*a*_ = .3 and set *ρ*_*e*_ to .2 or .5. In higher-dimensional settings (*k* > 10), we use a different structure that is based on having clusters within the *k* traits, with the correlation within clusters being higher than that between different clusters. We use non-overlapping clusters of 10 traits each, and within a given cluster, the correlation structure between traits is the same as in the case when *k* ≤ 10, where we also fix *ρ*_*a*_ = 0.3 and set *ρ*_*e*_ to .2 or .5. However, for traits in different clusters, we set the genetic correlation to 0.05 and the residual correlation to 0.01 (or 0.1) if *ρ*_*e*_ is 0.2 (or 0.5) within clusters. Additional details of Model I are in **Supplemental Methods**.

We also consider the simulation of a single binary trait using a liability threshold model, referred to as Model II, where the phenotype value *Y*_*i*_ for individual *i* is given by

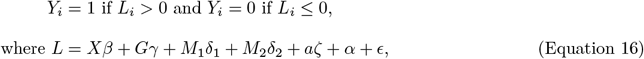

where *Y* is *n* ×1 with *Y*_*i*_ = 1 indicating that individual *i* is affected (i.e., a case subject) and *Y*_*i*_ = 0 that individual *i* is unaffected (i.e., a control subject); *L* is *n* ×1 with *L*_*i*_ being the underlying liability value for individual *i*; *X, G, M*_1_, *M*_2_, *a, β, γ, δ*_1_, *δ*_2_, *α*, and *ϵ* are as before, where now *k* = 1, which implies that *V*_*a*_ reduces to the scalar 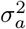 and *V*_*e*_ reduces to the scalar 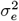.

#### Association test statistics considered

In the multivariate quantitative trait simulations, we consider two different test statistics, *T*_1_, which is defined in **Useful special cases I**, and *T*_2_, which is defined in **Useful special cases II**. Recall that test statistic *T*_1_ is based on a LMM that accounts for covariates and for covariance among the *k* traits, but ignores pedigree and population structure or only partially corrects for it by including some PCs as covariates. Test statistic *T*_2_ is based on a more elaborate LMM that accounts for covariates and for covariance among the *k* traits, and also accounts for pedigree and population structure among the sampled individuals by including an additive polygenic component of variance based on a GRM or kinship matrix. For *T*_2_, to obtain 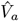 and 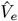 in the cases when *k* ≤ 5, we use GEMMA [47], and in the cases when *k* > 5, we use PHENIX [48]. For the binary trait simulations, we consider test statistic, *T*_3_, which is defined in **Use of JASPER to “robustify” a binary-trait, rare-variant association test**. *T*_3_ is a score test in a logistic mixed model that accounts for covariates and accommodates a weight matrix *W* for the rare variants, but ignores pedigree and population structure or only partially corrects for it by including some PCs as covariates. All three test statistics have the required form of Equation 1. The models used to derive two of the test statistics, *T*_1_ and *T*_3_, assume conditional independence of the trait values for different individuals under the null hypothesis given the covariates, which is not the case in our simulations (i.e., the phenotype models are misspecified as they ignore population structure). Hence, we consider including the top *l* PCs of the GRM *K* as covariates in the null models for *T*_1_ and *T*_3_, with *l* being either 1 or 10. Consequently, the matrix *X* used to fit *T*_1_ and *T*_3_ has either 4, 5, or 14 columns, and includes intercept, 3 observed covariates, and either 0, 1 or 10 PCs of the GRM, while the *X* matrix used to fit *T*_2_ has only 4 columns and does not include any PCs. In simulations, *T*_1_ and *T*_2_ are used with quantitative traits simulated from Model I, and *T*_3_ is used with binary traits simulated from Model II.

#### Comparison with other assessments of significance

We compare different methods to assess significance for *T*_1_, *T*_2_ and *T*_3_. For *T*_1_, we use for comparison MSKAT [10], which uses Davies’ method to analytically compute asymptotic p-values of *T*_1_. The resulting procedure is referred to as *T*_1_-asymp in what follows. For *T*_1_, we also use DKAT [28], which, like JASPER, uses a Pearson type III approximation to obtain p-values, but, unlike JASPER, assumes exchangeability of the rows of either *S*_*G*_ or *S*_*Y*_ under the null, which would hold if, for example, there was no population structure or if all the structure was captured by the PCs included as covariates (or if there was a very simple type of structure that preserved exchangeability). The resulting procedure is referred to as *T*_1_-perm in what follows. For *T*_2_, we compare our method to Multi-SKAT [12], which also uses Davies’ method to compute asymptotic p-values. For high dimensional traits (*k* ≥ 50), we were forced to reduce the number of replicates used to assess the type 1 error rate of Multi-SKAT to 3,000 and 1,200, for *k* = 50 and 100, respectively, owing to the computational burden of the Multi-SKAT method. The resulting procedure is referred to as *T*_2_-asymp. For *T*_3_, we compare our method to SKAT adapted to binary traits with a moment matching adjustment to compute asymptotic p-values [49]. The resulting procedure is referred to as *T*_3_-asymp. Lastly, for *T*_3_, we also compare our method to pedgene [32], which is a retrospective method of significance and is therefore robust to phenotype model misspecification, but which requires accurate estimation of *F*, the covariance matrix for the *m* genetic variants being tested. (In contrast, JASPER does not require estimation of *F* .) The resulting procedure is referred to as *T*_3_-retro-asymp. When we apply JASPER to assess significance of test statistic *T*_*i*_ (*i* = 1, 2, 3), we refer to the resulting procedure as *T*_*i*_-JASPER.

### Application to Gene Expression Data from the Framingham Heart Study

We apply JASPER to analyze gene expression data from the Framingham Heart Study (FHS) [50], which is a large, longitudinal, observational study that includes both unrelated individuals and individuals from multi-generation pedigrees. Our use of the FHS data was approved by the institutional review board of the Biological Sciences Division of the University of Chicago. We investigate the *cis* genetic architecture of gene expression in select biological pathways that have previously been associated with a wide range of disorders including inflammatory bowel disease, coronary heart disease, and type 2 diabetes. We focus on the Offspring Cohort and 11 gene pathways from the Kyoto Encyclopedia of Genes and Genomes (KEGG) database [51] (see Table 1). Gene expression profiling was carried out on a sample of 2,442 Offspring Cohort subjects using the Affymetrix Human Exon 1.0 ST Gene Chip platform, resulting in the profiling of 17,873 measured transcript clusters. After removing transcripts not mapping to RefSeq genes, 17,601 probes corresponding to 17,379 distinct genes remained. The expression values were normalized and residuals were obtained from a linear mixed model which included ten technical covariates (chip batch, RNA quality and 8 quality control metrics provided by Affymetrix APT program) [52]. We exclude *HLA* genes from the analysis, as these are the most polymorphic genes in the human genome and known hotspots for disease associations [53], and we also exclude genes not located on autosomes. Both age and sex are included as covariates in the analysis. Genotype data is obtained using the Affymetrix 500K mapping array set and we exclude from the analysis individuals who have (1) completeness < 96%; (2) missingness > 10% in any of the genetic regions analyzed; or (3) empirical self-kinship coefficient greater than > .525 (i.e., empirical inbreeding coefficient > .05). We also exclude a few individuals whose off-diagonal empirical kinship coefficient values appear inconsistent with the given pedigree information. This results in 1,894 individuals with covariate, genotype and expression data that are included in the analysis. For each gene in a given KEGG pathway, we extract all the polymorphic sites on the Affymetrix 500K chip that are within 500kb of the gene. We exclude sites that have (1) call rate ≤ 96%, or (2) Mendelian error rate > 2%. For the remaining sites, we impute any missing genotypes using IMPUTE2 [54]. This yields a total of 182,447 SNPs mapped to 1,194 genes across the 11 KEGG pathways, with a median number of *cis*-SNPs for any given gene of 152. We conduct 3 sets of association analyses in which we evaluate the association between *cis*-SNPs of a given gene and (1) the expression values of the gene; (2) the expression values of all genes in the pathway excluding the gene; (3) the expression values of all genes in the pathway including the gene. The three sets of analyses are referred to as “SINGLE” (for single gene), “LOGO” (for leave one gene out), and “ALL” (for all genes in the pathway), respectively.

**Table 1:** List of KEGG pathways analyzed with the FHS data.

| Pathway name | KEGG ID | Number of Genes<sup>a</sup> |
| --- | --- | --- |
| Antigen processing and presentation | hsa04612 | 42 |
| Cytokine-cytokine receptor interaction | hsa04060 | 247 |
| JAK-STAT signaling pathway | hsa04630 | 141 |
| Inflammatory bowel disease | hsa05321 | 49 |
| Autophagy | hsa04140 | 106 |
| cGMP-PKG signaling pathway | hsa04022 | 151 |
| T cell receptor signaling pathway | hsa04660 | 94 |
| Neurotrophin signaling pathway | hsa04722 | 111 |
| Wnt signaling pathway | hsa04310 | 144 |
| PPAR signaling pathway | hsa03320 | 66 |
| Notch signaling pathway | hsa04330 | 43 |

<sup>a</sup> Excludes *HLA* genes, genes not on autosomes and genes that do not have transcripts in the FHS data

## Results

### Type 1 error and power studies

In simulations, we evaluate the JASPER method for assessing significance of tests of single- and multitrait association with sets of multiple genetic variants when there is population and/or pedigree structure in the data, taking into account relevant covariates. Specifically, we evaluate the ability of JASPER to correctly control type 1 error and maintain high power, even when the statistic to which it is applied either ignores the population and pedigree structure or only partially accounts for it, in which case most other methods to assess significance are expected to fail. We also evaluate the ability of JASPER to provide correct type 1 error control and maintain high power when assessing significance with high-dimensional traits, when asymptotic assessments of significance can break down, even if population and pedigree structure is appropriately accounted for. Finally, we evaluate the ability of JASPER to provide correct type 1 error control and maintain high power when assessing significance of a binary trait with a large number of rare variants, when a retrospective, asymptotic assessment of significance can break down. In addition, we assess robustness of JASPER to misspecification of the phenotype model.

### JASPER can “Robustify” single- or multitrait test statistics that do not fully correct for sample structure

First, we consider a setting in which association between a single quantitative trait and a set of 50 variants is tested, in a sample with both population and pedigree structure. We consider the statistic *T*_1_ which either ignores the population and pedigree structure (when 0 PCs are included as covariates) or only partially corrects for it (when 1 or 10 PCs are included as covariates). In Table 2, in the *T*_1_-JASPER column, we can see that use of JASPER to assess significance of *T*_1_ results in correct type 1 error in all settings, showing that JASPER is able to successfully “robustify” *T*_1_ to make it valid in a context of population and pedigree structure. In contrast, use of either an asymptotic assessment of significance or an approximation to a naive permutation test, which both ignore the sample structure, lead to drastically inflated type 1 error (*T*_1_-asymp and *T*_1_-perm columns of Table 2).

**Table 2:** Empirical Type 1 Error for Association Between a Single Quantitative Trait and a SNP Set, with 2 Sub-populations and Relatedness.

| # PCs<sup>b</sup> | Nominal Level | Empirical Type 1 Error<sup>a</sup> of |  |  |  |  |
| --- | --- | --- | --- | --- | --- | --- |
| | | $T_1$-JASPER | $T_1$-perm | $T_1$-asyp | $T_2$-JASPER | $T_2$-asyp |
| 0 | 0.05 | 0.0500 | 1.0000* | 1.0000* | 0.0480 | 0.0428* |
| 0 | 0.005 | 0.0043 | 1.0000* | 1.0000* | 0.0045 | 0.0035* |
| 1 | 0.05 | 0.0503 | 0.6761* | 0.6893* | - | - |
| 1 | 0.005 | 0.0051 | 0.3615* | 0.3652* | - | - |
| 10 | 0.05 | 0.0514 | 0.5370* | 0.6360* | - | - |
| 10 | 0.005 | 0.0054 | 0.2344* | 0.3125* | - | - |

The quantitative trait is generated from Model I.
<sup>a</sup> Empirical type 1 error is based on 30,000 replicates. A star indicates a type 1 error rate that is significantly different from the nominal level, using the z-test at level .01. The .01-level acceptance interval for the test when the nominal rate is .05 is (.0458,.0532), and the .01-level acceptance interval when the nominal rate is .005 is (.00395,.00605).
<sup>b</sup> # PCs denotes the number of top principal components of the estimated GRM included as covariates in the test statistic $T_1$ .

JASPER can be applied equally well to binary traits. We obtain similar results (Table 3) in a setting in which association between a single binary trait and a set of 50 variants is tested, in a sample with both population and pedigree structure, using the binary trait test statistic *T*_3_, which only partially corrects for sample structure by including either 1 or 10 PCs as covariates. We again see that the type 1 error of JASPER is correct in all settings, while type 1 error can be drastically inflated by standard methods for assessing significance (“*T*_3_-asymp” column of Table 3) when the test statistic fails to adequately capture population structure.

**Table 3:**
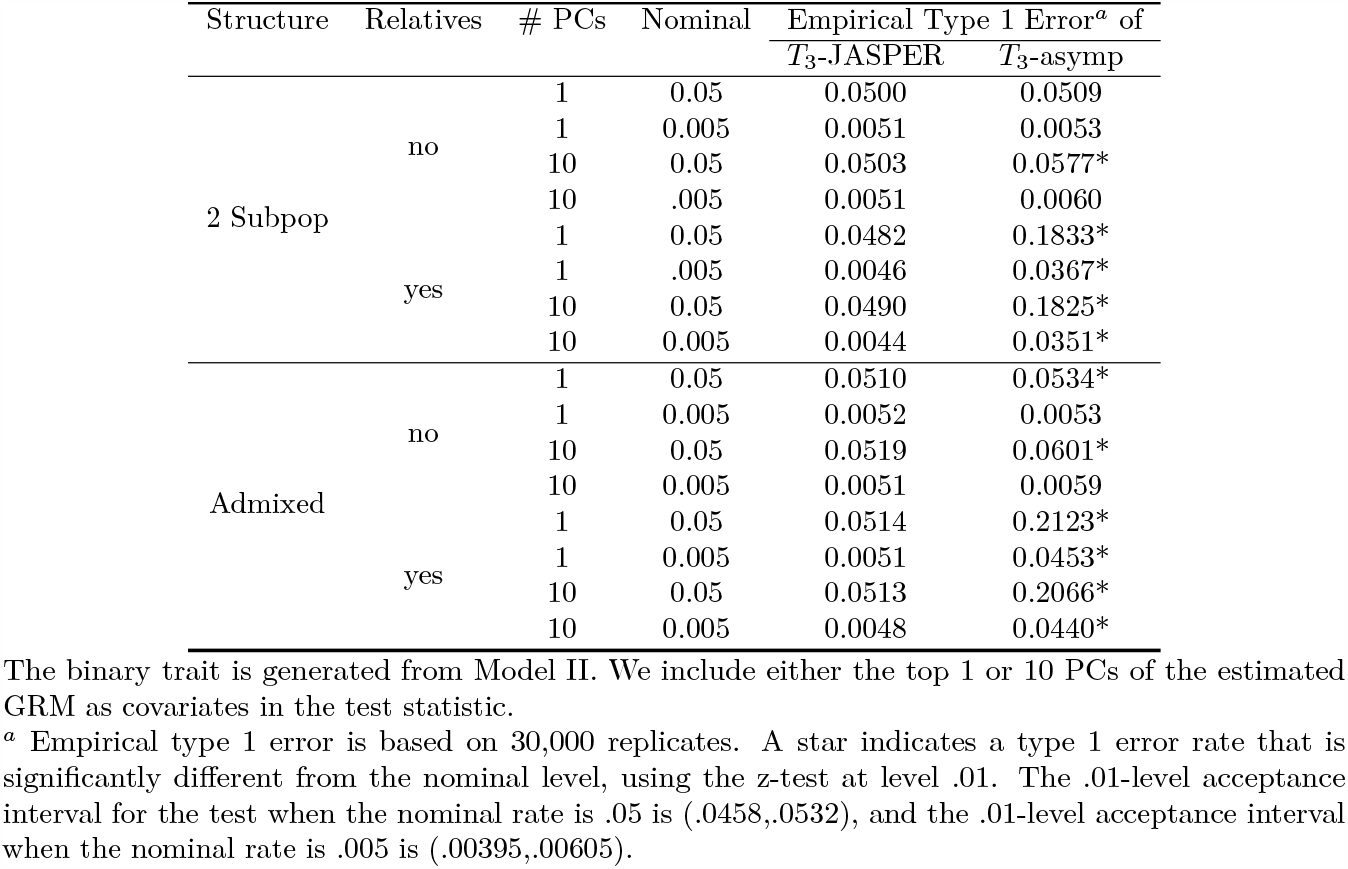
Empirical Type 1 Error for Association Between a Binary Trait and a SNP set, with Population and Pedigree Structure.

| Structure | Relatives | # PCs | Nominal | Empirical Type 1 Error<sup>a</sup> of |  |
| --- | --- | --- | --- | --- | --- |
| | | | | $T_3$-JASPER | $T_3$-asyp |
| 2 Subpop | no | 1 | 0.05 | 0.0500 | 0.0509 |
|  |  | 1 | 0.005 | 0.0051 | 0.0053 |
|  |  | 10 | 0.05 | 0.0503 | 0.0577* |
|  |  | 10 | .005 | 0.0051 | 0.0060 |
|  | yes | 1 | 0.05 | 0.0482 | 0.1833* |
|  |  | 1 | .005 | 0.0046 | 0.0367* |
|  |  | 10 | 0.05 | 0.0490 | 0.1825* |
|  |  | 10 | 0.005 | 0.0044 | 0.0351* |
| Admixed | no | 1 | 0.05 | 0.0510 | 0.0534* |
|  |  | 1 | 0.005 | 0.0052 | 0.0053 |
|  |  | 10 | 0.05 | 0.0519 | 0.0601* |
|  |  | 10 | 0.005 | 0.0051 | 0.0059 |
|  | yes | 1 | 0.05 | 0.0514 | 0.2123* |
|  |  | 1 | 0.005 | 0.0051 | 0.0453* |
|  |  | 10 | 0.05 | 0.0513 | 0.2066* |
|  |  | 10 | 0.005 | 0.0048 | 0.0440* |

The binary trait is generated from Model II. We include either the top 1 or 10 PCs of the estimated GRM as covariates in the test statistic.
<sup>a</sup> Empirical type 1 error is based on 30,000 replicates. A star indicates a type 1 error rate that is significantly different from the nominal level, using the z-test at level .01. The .01-level acceptance interval for the test when the nominal rate is .05 is (.0458,.0532), and the .01-level acceptance interval when the nominal rate is .005 is (.00395,.00605).

For multitrait mapping, where we test association between a set of *k* traits and a set of 50 SNPs in a sample with both population and pedigree structure, JASPER is again able to robustify *T*_1_. In the *T*_1_-JASPER column of Table 4, we can see that the type 1 error of JASPER is correct in all settings.

**Table 4:** Empirical Type 1 Error for Multitrait Association with a SNP Set, with 2 Sub-populations and Relatedness.

| # Traits $k$ | $\rho_e^b$ | Nominal Level | Empirical Type 1 Error<sup>a</sup> of | | |
| --- | --- | --- | --- | --- | --- |
| | | | $T_1$-JASPER<sup>c</sup> (SE) | $T_2$-JASPER (SE) | $T_2$-asympt (SE) |
| 2 | 0.20 | 0.05 | 0.0491 (.0012) | 0.0485 (.0012) | 0.0424* (.0012) |
|  |  | 0.005 | 0.0054 (.0004) | 0.0048 (.0004) | 0.0036* (.0003) |
|  | 0.50 | 0.05 | 0.0511 (.0012) | 0.0509 (.0012) | 0.0460* (.0012) |
|  |  | 0.005 | 0.0048 (.0004) | 0.0052 (.0004) | 0.0043 (.0004) |
| 5 | 0.20 | 0.05 | 0.0516 (.0012) | 0.0506 (.0012) | 0.0438* (.0012) |
|  |  | 0.005 | 0.0056 (.0004) | 0.0058 (.0004) | 0.0041 (.0004) |
|  | 0.50 | 0.05 | 0.0509 (.0012) | 0.0494 (.0012) | 0.0440* (.0012) |
|  |  | 0.005 | 0.0049 (.0004) | 0.0050 (.0004) | 0.0040 (.0004) |
| 50 | 0.20 | 0.05 | 0.0492 (.0012) | 0.0510 (.0012) | 0.0003* (.0003) |
|  |  | 0.005 | 0.0049 (.0004) | 0.0045 (.0004) | 0.0003* (.0003) |
|  | 0.50 | 0.05 | 0.0511 (.0012) | 0.0527 (.0012) | 0.0003* (.0003) |
|  |  | 0.005 | 0.0051 (.0004) | 0.0058 (.0004) | 0.0003* (.0003) |
| 100 | 0.20 | 0.05 | 0.0525 (.0012) | 0.0501 (.0012) | 0.0008* (.0008) |
|  |  | 0.005 | 0.0055 (.0004) | 0.0050 (.0004) | 0.0008 (.0008) |
|  | 0.50 | 0.05 | 0.0526 (.0012) | 0.0496 (.0012) | 0.0008* (.0008) |
|  |  | 0.005 | 0.0060 (.0004) | 0.0048 (.0004) | 0.0008 (.0008) |

The $k$ traits are generated from Model I.
<sup>a</sup> The empirical type 1 error of $T_1$ -JASPER and $T_2$ -JASPER is based on 30,000 replicates in all cases. The empirical type 1 error of $T_2$ -asympt is based on 30,000 replicates for $k = 2$ or 5, 3,000 replicates for $k = 50$ , and 1,200 replicates for $k = 100$ . A star indicates a type 1 error rate that is significantly different from the nominal level, using the z-test at level .01 if the expected count is at least 10 or an exact binomial test at level .01 if the expected count is less than 10.
<sup>b</sup> $\rho_e$ is the residual correlation between all traits (for $k < 50$ ) or between traits within clusters of size 10 (for $k \geq 50$ ).
<sup>c</sup> The top 10 PCs are included as covariates in the trait model for $T_1$ -JASPER.

Retrospective, asymptotic assessment of significance [16, 30–34] is a different approach that is also able to “robustify” a test statistic that either ignores or does not fully capture population and pedigree structure. In Table 5, we compare JASPER with a retrospective, asymptotic assessment of significance for testing a binary trait with multiple rare variants [32], which we call “retro-asymp” in Table 5, where we use both methods to assess significance of *T*_3_ in a sample with pedigree structure. Although *T*_3_ does not adequately account for the pedigree structure, we can see in Table 5 that both JASPER and retro-asymp are able to “robustify” the test statistic, in the sense that neither shows inflated type 1 error. However, as the variants become rarer, retro-asymp becomes highly conservative, while JASPER maintains correct type 1 error. In Table 6, we can see that JASPER also has consistently higher power than retro-asymp. We believe that the conservativeness of retro-asymp is due to the challenge of accurately estimating *F*, the covariance matrix of the rare variants, which is required for retro-asymp. In contrast, JASPER does not require knowledge of *F* and retains accurate type 1 error and high power even as the variants become rarer.

**Table 5:**
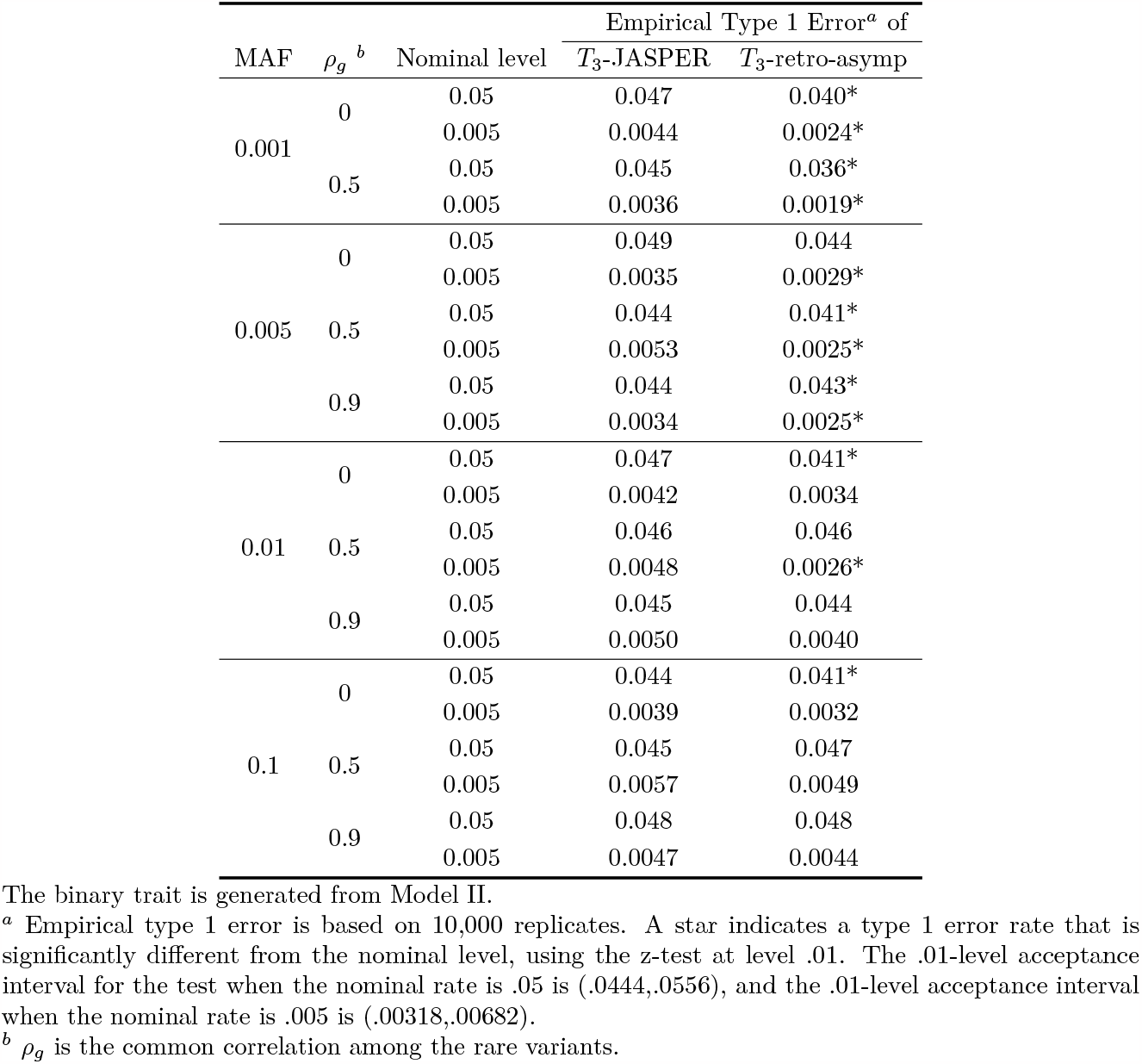
Empirical Type 1 Error for Association Between a Binary Trait and a Set of Rare Variants, with Related Individuals.

| MAF | $\rho_g^b$ | Nominal level | Empirical Type 1 Error<sup>a</sup> of | |
| --- | --- | --- | --- | --- |
| | | | $T_3$-JASPER | $T_3$-retro-asymp |
| 0.001 | 0 | 0.05 | 0.047 | 0.040* |
|  |  | 0.005 | 0.0044 | 0.0024* |
|  | 0.5 | 0.05 | 0.045 | 0.036* |
|  |  | 0.005 | 0.0036 | 0.0019* |
| 0.005 | 0 | 0.05 | 0.049 | 0.044 |
|  |  | 0.005 | 0.0035 | 0.0029* |
|  | 0.5 | 0.05 | 0.044 | 0.041* |
|  |  | 0.005 | 0.0053 | 0.0025* |
|  | 0.9 | 0.05 | 0.044 | 0.043* |
|  |  | 0.005 | 0.0034 | 0.0025* |
| 0.01 | 0 | 0.05 | 0.047 | 0.041* |
|  |  | 0.005 | 0.0042 | 0.0034 |
|  | 0.5 | 0.05 | 0.046 | 0.046 |
|  |  | 0.005 | 0.0048 | 0.0026* |
|  | 0.9 | 0.05 | 0.045 | 0.044 |
|  |  | 0.005 | 0.0050 | 0.0040 |
| 0.1 | 0 | 0.05 | 0.044 | 0.041* |
|  |  | 0.005 | 0.0039 | 0.0032 |
|  | 0.5 | 0.05 | 0.045 | 0.047 |
|  |  | 0.005 | 0.0057 | 0.0049 |
|  | 0.9 | 0.05 | 0.048 | 0.048 |
|  |  | 0.005 | 0.0047 | 0.0044 |

The binary trait is generated from Model II.
<sup>a</sup> Empirical type 1 error is based on 10,000 replicates. A star indicates a type 1 error rate that is significantly different from the nominal level, using the z-test at level .01. The .01-level acceptance interval for the test when the nominal rate is .05 is (.0444,.0556), and the .01-level acceptance interval when the nominal rate is .005 is (.00318,.00682).
<sup>b</sup> $\rho_g$ is the common correlation among the rare variants.

**Table 6:** Empirical Power for Association Between a Binary Trait and a Set of Rare Variants, with Pedigree Structure.

| MAF | $\rho_g$<sup>b</sup> | Nominal level | Empirical Power<sup>a</sup> (SE) of | |
| --- | --- | --- | --- | --- |
| | | | $T_3$-JASPER | $T_3$-retro-asymp |
| 0.001 | 0 | 0.001 | 0.290 (.014) | 0.251 (.014) |
|  |  | 0.005 | 0.408 (.016) | 0.389 (.015) |
|  | 0.5 | 0.001 | 0.288 (.014) | 0.256 (.014) |
|  |  | 0.005 | 0.395 (.015) | 0.378 (.015) |
|  | 0.9 | 0.001 | 0.266 (.014) | 0.229 (.013) |
|  |  | 0.005 | 0.372 (.015) | 0.350 (.015) |
| 0.005 | 0 | 0.001 | 0.242 (.014) | 0.198 (.013) |
|  |  | 0.005 | 0.408 (.016) | 0.383 (.015) |
|  | 0.5 | 0.001 | 0.270 (.014) | 0.226 (.013) |
|  |  | 0.005 | 0.464 (.016) | 0.414 (.016) |
|  | 0.9 | 0.001 | 0.278 (.014) | 0.233 (.013) |
|  |  | 0.005 | 0.465 (.016) | 0.428 (.016) |

The binary trait is generated from Model II.
<sup>a</sup> Empirical power is based on 1,000 replicates.
<sup>b</sup> $\rho_g$ is the common correlation among the rare variants.

### JASPER can work better than asymptotic methods regardless of the number of traits

Even when the test statistic appropriately takes into account the population and pedigree structure in the sample, JASPER can still have better type 1 error and power than asymptotic methods. This can be seen in our simulation studies that use the test statistic *T*_2_. Even with only one quantitative trait, Table 2 shows that *T*_2_-JASPER has correct type 1 error while *T*_2_-asymp is somewhat conservative. In Table 4, we can see that for a set of traits ranging in size from *k* = 2 to *k* = 100, *T*_2_-JASPER provides correct type 1 error, while *T*_2_-asymp ranges from somewhat conservative for *k* = 2 to exceedingly conservative for *k* = 100. This results in large power differences between the two methods (Table 7), with *T*_2_-JASPER providing consistently higher power, and the power of *T*_2_-asymp dropping to near zero for *k* = 100. The collapse of power of *T*_2_-asymp for larger *k* might be due to the estimation error incurred by estimating the two *k* ×*k* covariance matrices between traits (*V*_*a*_ and *V*_*e*_ in the null model of Equation 12), where assessment of significance for *T*_2_-asymp presumably relies on reasonably accurate estimation of *V*_*a*_ and *V*_*e*_. However, *T*_2_-JASPER is still able to maintain correct type 1 error and power, despite using the same estimation approach to construct the *T*_2_ statistic, because JASPER’s assessment of significance does not rely on accurate estimation of *V*_*a*_ or *V*_*e*_. Overall, we find that, in addition to providing robustness in type 1 error control, assessing significance using JASPER can greatly increase power compared to an asymptotic assessment of significance, particularly when the number of traits analyzed is large.

**Table 7:**
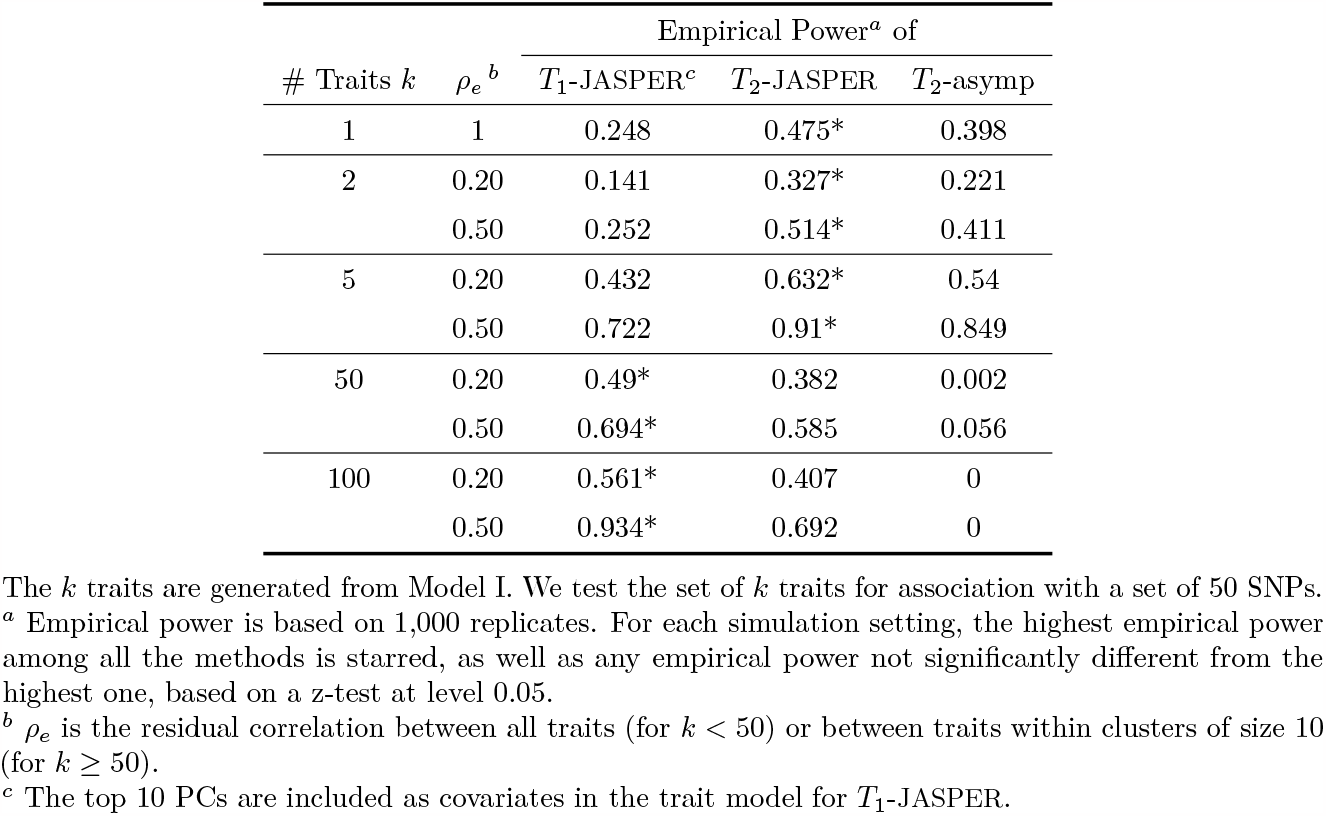
Empirical Power for Multitrait Association with a SNP set, with 2 Sub-populations and Relatedness.

Comparison of the two underlying test statistics *T*_1_ and *T*_2_ is somewhat beyond the scope of this work (which is instead focused on comparison of methods of assessment of significance). Nonetheless, it is interesting to note that for a small number of traits, *T*_2_-JASPER is able to increase power over *T*_1_-JASPER by including detailed population and pedigree structure information in a GRM, while *T*_1_-JASPER only includes the partial information of 10 PCs of the GRM (comparison of power of *T*_1_-JASPER and *T*_2_-JASPER in Table 7 for *k* ≤ 5). However, as the number of traits grows large, the comparison reverses, and *T*_1_-JASPER has higher power than *T*_2_-JASPER for *k* = 50 or 100 in Table 7. The presumably reflects the trade-off in power between including additional population structure information, which would be expected to increase power, vs. estimating an additional (*k* + 1)*k/*2 parameters in *T*_2_ (due to *V*_*a*_ in Equation 12), which could be expected to decrease power as *k* grows large. An advantage of using JASPER to assess significance is that one is free to choose between test statistics (e.g., *T*_1_ and *T*_2_) purely based on power, without worrying about the fact that one statistic (*T*_2_) fully corrects for population structure and the other (*T*_1_) does not, because JASPER is effective at controlling the type 1 error, regardless.

The simulation studies also show that JASPER is robust to various sources of misspecification in the trait model including (1) unobserved covariates (i.e., the two unobserved causal genes that were included in every simulation setting but left out of the fitted model in every case); (2) use of the wrong link function for a binary trait (*T*_3_-JASPER in Tables 3, 5 and 6, where a logistic model is fit to a trait simulated under a liability threshold model); as well as to (3) misspecification of the population and pedigree structure in the fitted model (*T*_1_-JASPER in Tables 2, 4, and 7 and *T*_3_-JASPER in Tables 3, 5 and 6). This is not surprising given that the theoretical validity of JASPER essentially does not depend on the phenotype model at all.

Overall, these results demonstrate the superiority of JASPER over the asymptotic approximations, both prospective and retrospective, and the Pearson type III approximation assuming unrelated samples, in terms of type 1 error control and statistical power for association testing in structured samples.

### Analysis of Expression Data from Framingham Heart Study

We perform single- and multitrait analysis of FHS expression data, using the SINGLE, LOGO and ALL approaches described in **Application to Gene Expression Data from the Framingham Heart Study**. For all three approaches, analysis is performed with each of *T*_1_-JASPER, *T*_2_-JASPER and *T*_1_-asymp. The statistic *T*_2_-asymp is only used in the SINGLE analysis, which corresponds to a single gene’s expression values (i.e., *k* = 1) being tested with its *cis*-SNPs, as it is computationally challenging to use it for the other two analyses which involve a much larger number of traits (*k* > 100 in the majority of cases). For all statistics, we include age and sex as covariates, and for *T*_1_-asymp and *T*_1_-JASPER, we include the top 10 PCs of the estimated GRM as covariates in the trait model as well.

In the simulation studies, we showed that by properly accounting for population structure JASPER can improve both type 1 error and power. In the data analysis, examples of both effects can be seen in Figure 1 Panels A and B, which compare p-values from 3 methods for testing association between a set of *cis*-SNPs for a given gene and the expression levels of the genes in the pathway of the given gene. In Panel A, we can see that *T*_1_-asymp has consistently inflated p-values compared to *T*_1_-JASPER and *T*_2_-JASPER for the bulk of the p-values, which is what we expect based on simulation studies showing that both JASPER methods maintained correct type 1 error by properly accounting for population structure, while *T*_1_-asymp had inflated type 1 error when population structure was not accounted for at all (0 PCs included) or only partially accounted for (1 or 10 PCs included). At the same time, in Panel B, we can see that the *T*_1_-JASPER method has consistently more significant results than *T*_1_-asymp for the strongest signals, which is consistent with a power improvement due to appropriately accounting for population structure. Similarly, for association of a set of SNPs with a single trait, Table 8 shows that additional detections of significant SNPs are made when the test that properly accounts for population structure (*T*_2_-JASPER) is used compared to a test that only partially accounts for population structure (*T*_1_-asymp).

**Table 8:** Association detected between expression of a single gene and its set of *cis*-SNPs in the Framingham data.

|  |  |
| --- | --- |
| Genes significant by both $T_2$-JASPER and $T_1$-asyp | 165 |
| Genes significant by $T_2$-JASPER only | 27 |
| Genes significant by $T_1$-asyp only | 0 |

**Figure 1:**
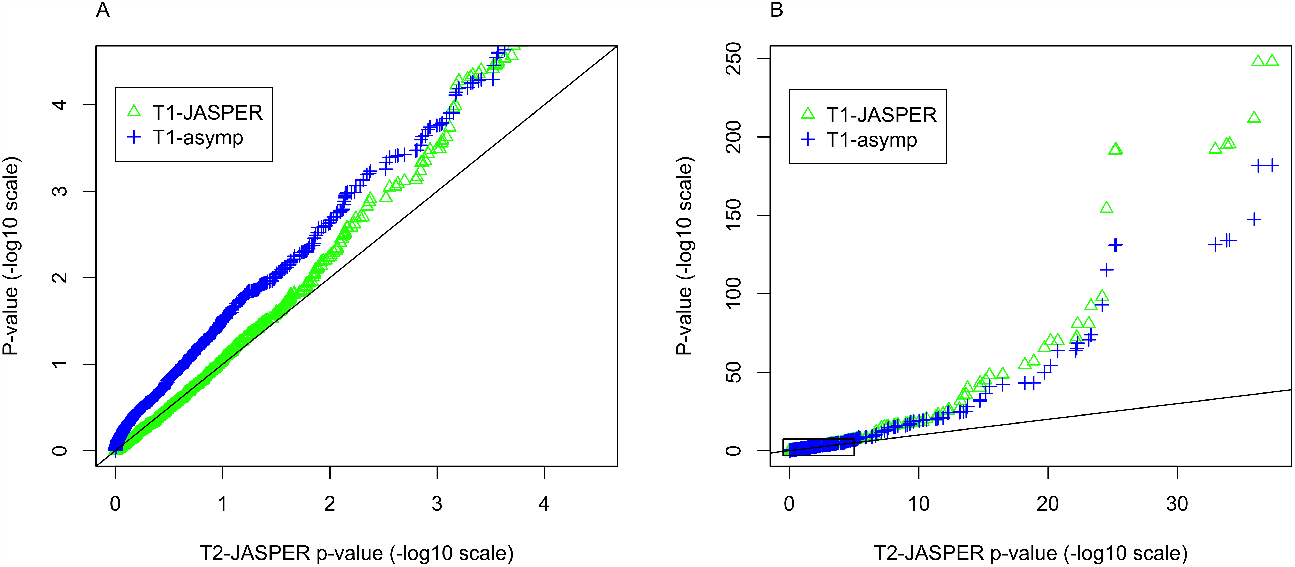
Comparison of P-values for Multitrait Analysis in Framingham Data when Population Structure is or is not Fully Accounted for. QQ plot comparing the ordered p-values on the − log_10_ scale for three different association testing methods. In Panel A, only the large p-values are plotted, and in Panel B, all the p-values are plotted, where the small rectangle in Panel B indicates the part of the plot that is shown in Panel A.

Table 9 compares the SNP sets that are detected by a single-trait analysis to those detected by a multitrait analysis. We use *T*_2_-JASPER for the single trait analysis and *T*_1_-JASPER for the multitrait analysis because the simulation studies indicated that these were the most powerful test statistics in those contexts, among those we considered. In Table 9, we can see that while the single- and multitrait analysis have substantial overlap in the SNP sets detected, there are also many SNP sets detected by only one or the other, indicating that the two analyses are to some extent complementary and provide different types of information on association.

**Table 9:** Number of SNP sets for which significant association was detected in either a multitrait or single-trait analysis in the Framingham data.

|  |  |
| --- | --- |
| SNP sets significant in both single- and multitrait analyses | 79 |
| SNP sets significant in single-trait analysis only | 114 |
| SNP sets significant in multitrait analysis only | 27 |

Tables S1-S3 report the gene regions for which the p-value for *T*_2_-JASPER in the ALL analysis is below 4.2 ×10^−5^, which corresponds to a Bonferroni correction based on the 1,194 genes being tested across the 11 pathways. We find many genes with highly significant p-values in both the SINGLE and ALL analyses with *T*_1_-JASPER and *T*_2_-JASPER, such as *KLRC3* in pathway hsa04612, *SCP2* in pathway hsa03320, *IL18RAP* in pathway hsa04060, and *DAPK1* in pathway hsa04140. The results from the LOGO analysis help us elucidate whether the signal detected in the ALL analysis is driven mostly by the effect of a set of *cis*-SNPs on the gene it is nearest to or if it is also driven by the association of the *cis*-SNPs with the expression of other genes in the same pathway. Among the genes where both the SINGLE and ALL p-values were significant, only a few genes also had significant p-values in the LOGO analysis: *KLRC1, KLRD1* and *KLRC3* in pathway hsa04612, *IL1R2, IL1RL1, IL18R1, IL18RAP, CCR2, CCR1*, and *CCR3* in pathway hsa04060, *IL18RAP* and *IL18R1* in pathway hsa05321, and *CALML4* in pathway hsa04722. Eight of these genes have previously been associated with a disease that has been linked to the corresponding KEGG pathway [55]. For the remaining five genes, the analysis suggests that the corresponding KEGG pathway is a candidate pathway for association with the diseases the genes have been previously linked to. For example, the gene *IL1RL1* in the ‘Cytokine-cytokine receptor interaction’ pathway has been previously associated with asthma [56, 57], and *CCR1* in the same pathway has been previously associated with Behcet’s disease [58, 59], a form of vasculitis, but neither of the two diseases has previously been associated with the KEGG pathway.

One could also look at genes that have a significant p-value in the LOGO analysis but not in the SINGLE analysis, as this would suggest that the *cis*-SNPs of the gene have *trans* effects in the pathway (i.e., affecting genes other than the nearest one). Examples of such genes are *HSPA1A* in pathway hsa04612, *CPT2* in pathway hsa03320, *NUMB* in pathway hsa04330, *IL1RL2* and *IL1R1* in pathway hsa04060, and *IL23R* in pathway hsa05321. Six out of twelve of such genes have been previously associated with a disease linked to the KEGG pathway. The remaining genes could be candidate genes to study in a GWA analysis with the diseases linked to the corresponding KEGG pathway. Overall, seven out of the eleven KEGG pathways analyzed have, among the top 3 signals, genes that were previously associated with diseases linked to the pathway.

### Computation time of JASPER

The main computational burden of JASPER lies in the eigen-decomposition of the *n* ×*n* matrix that is needed to correct for population structure. However, the test statistic being used may already require the eigen-decomposition of the same matrix, in which case the results of that decomposition could be re-used by JASPER. Using a single processor (at 3.5GHz) of Intel Xeon CPU E5-2637 v3, we compare the runtime for assessing significance for a single gene region in the FHS data using JASPER to that for an asymptotic approximation. We pick the largest KEGG pathway ‘Cytokine-cytokine receptor interaction’ (hsa04060) containing 247 genes, and test the association between a gene region and gene expression values at *k* genes in the pathway. We pick 3 gene regions corresponding to 51 *cis*-SNPs, 100 *cis*-SNPs, and 152 *cis*-SNPs. For each region, we sample at random *k* genes in the pathway for which we obtain the expression values, and the statistic *T*_2_ is used for association testing. We exclude the time taken to obtain the GRM eigen-decomposition (which is required in both methods). Results are shown in Figure 2, where we see that the computation time for JASPER is approximately constant as the number of tested variants *m* and the number of traits *k* increase. This is because the computations for the JASPER method are based on two *n* ×*n* matrices. The run time for the asymptotic method increases dramatically as either *m* or *k* increases, because it relies on the eigen-decomposition of *mk* ×*mk* matrices. This demonstrates the computational efficiency of JASPER in high dimensional settings.

**Figure 2:**
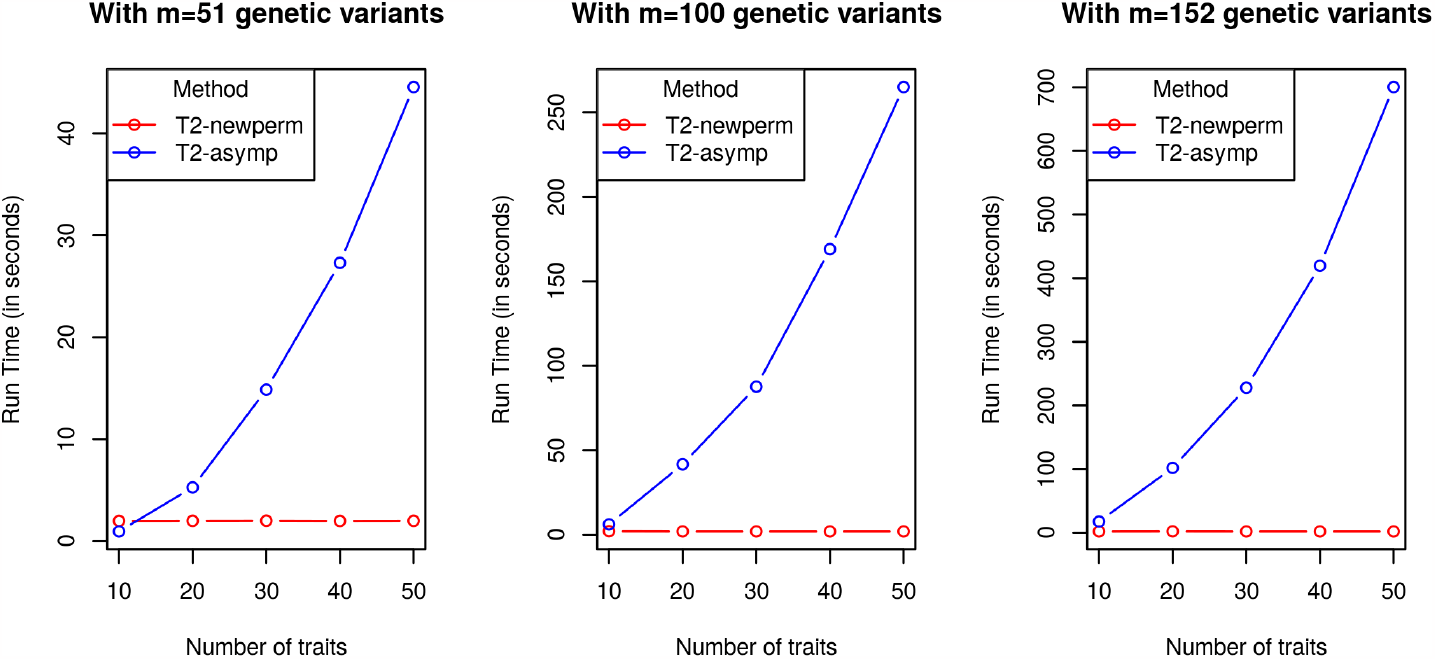
Run Time Comparison for JASPER and a Large Sample-based Method, in the FHS Data Using the Test Statistic *T*_2_. The JASPER results are denoted by *T*_2_-newperm, while the large sample-based results are denoted by *T*_2_-asymp.

## Discussion

We have developed JASPER, a fast, powerful and robust method for assessing significance of genetic association between a set of traits and a set of genetic variants, in samples that have population sub-structure, admixture and/or relatedness. JASPER is a versatile tool with many possible applications, including testing genetic association for multiple disease traits, expression traits, image-derived traits and microbiome abundances. JASPER is applicable to a wide range of association test statistics, including kernel-based (“KAT”-type) tests with essentially arbitrary phenotype kernels, and it is well-suited to handling high-dimensional traits. JASPER allows for incorporation of covariates, and it corrects for population structure by making using of a GRM or kinship matrix. Simulation studies show that JASPER results in higher power, better type 1 error control, and faster computation compared to existing methods for assessing significance in structured samples, with dramatically higher power and faster computation time for JASPER when a large number of traits is analyzed simultaneously.

We demonstrate the of use JASPER to (1) derive a fast, robust and powerful assessment of significance for a multitrait, multi-marker association test that allows for population structure, in a high dimensional setting in which an asymptotic assessment of significance breaks down; (2) start with a multitrait, multi-marker association test that does not adequately account for population structure, and “robustify” it, so that it properly accounts for population structure and results in a powerful test; and, similarly, (3) “robustify” a rare variant binary trait association test that does not adequately account for population structure, so that it becomes valid and powerful in a sample with population structure. In simulations, we show its effectiveness for all of these tasks, demonstrating that JASPER has the robustness advantages of a permutation test while being applicable to structured samples like an asymptotic assessment of significance. At the same time, JASPER is computationally faster than both of these other approaches.

In analysis of the FHS data, we find that the JASPER approach provides both better type 1 error control and higher power than the analytical assessment of significance for detecting association between a set of SNPs in a gene and the expression levels of a set of genes in a KEGG pathway. We find many significant associations, including 25 that indicate trans effects in which cis SNPs for one gene have significant association with a different set of genes in the same KEGG pathway. Of these 25, 14 have previously been associated with a disease linked to the KEGG pathway, and the remaining 11 associations have not previously been reported and could be candidate genes for future GWAS analysis with the diseases linked to the corresponding KEGG pathway. Our results demonstrate the promise of JASPER for powerful multitrait analysis in structured samples.

Because of its retrospective approach to assessing significance, JASPER is inherently robust to phenotype model misspecification. We show theoretically (Proposition 1 in **Appendix A**) that it is valid for kernel-based association statistics with essentially arbitrary phenotype kernels. In simulations, we demonstrate robustness to (1) unobserved covariates, (2) incorrect link function for a binary phenotype, and (3) failure to adequately model pedigree or population sub-structure. For the same reasons, JASPER would to be robust to phenotype-based ascertainment, which is basically just another type of phenotype model misspecification. Both Proposition 1 and the simulations also show that it is not necessary to have accurate estimation (or, in fact, any estimation at all) of either trait correlation or correlation among the tested genetic variants, in order for JASPER to have correct type 1 error (though this information, if available, can be easily incorporated, and in some cases can increase power). Although binary traits are not the primary focus of this paper, we have shown that JASPER has the potential to address a particularly challenging problem with binary traits, namely, that computational constraints have previously made it difficult to adequately incorporate correction for population and sample structure, while maintaining high power by modeling the special mean-variance structure of a binary trait [44], particularly in a multitrait, multi-variant context.

JASPER also works well for single- or multitrait association testing in samples in which there is no sample structure, in which case the decorrelation step described in **Decorrelation of** *S*_*G*_ **in JASPER** can be skipped, and the JASPER method would be equivalent to DKAT [28]. In principle, JASPER can be applied to more general association testing problems, not just phenotype-genotype association testing, provided that either (1) there is some way to decorrelate the rows and columns of one of the two kernel matrices or (2) there is no sample structure among the individuals.

Although we have emphasized multitrait mapping, we have also shown in simulations that JASPER can be useful for association testing of univariate traits as well, with multiple rare or common variants, or even just one variant. In that setting, asymptotic assessments of significance tend to be overly conservative, and our simulations show that JASPER provides better type 1 error control and higher power. In this paper we have exclusively focused on decorrelation of the rows and columns of *S*_*G*_ in using JASPER, but in principle we could instead decorrelate the rows and columns of *S*_*Y*_. In the LMM of Equation 12, it would not be obvious how to do this unless all the traits had the same heritability (or there were no sample structure). However, for a univariate trait, it would be straightforward to use the modeled variance structure of *Y*, with estimated parameters, to decorrelate the rows and columns of *S*_*Y*_ to provide a different way to use JASPER to assess significance. This approach would of course be sensitive to phenotype model misspecification, like the asymptotic approaches. However, for a correctly specified phenotype model, this alternative use of JASPER would still be expected to provide more accurate type 1 error and higher power than asymptotic approaches because the asymptotic approaches tend to be overly conservative.

In single- or multitrait genetic association analysis with a set of genetic variants, it is typical that the true phenotype model is unknown. Therefore, in kernel-based genetic association tests one approach to trying to maintain high power across a range of possible alternative models is to create an omnibus test statistic by combining multiple kernels, either through a weighted linear combination of kernels or adaptively, by taking the one with the smallest p-value in the observed data. Zhan et al. [28] have assessed both omnibus approaches by simulation and shown that the weighted linear combination approach is more powerful. The JASPER method can be directly applied to the weighted linear combination approach by simply taking that weighted linear combination to be *S*_*Y*_ (though it still requires a linear or weighted linear genotype kernel for *S*_*G*_). For a test statistic that involves selecting the kernel with the smallest p-value in the observed data, the JASPER method we describe is not directly applicable, and it could be of possible future interest to try to extend the method to that case. Another direction for possible future work could be to consider refinements to the estimator of the row covariance matrix of *G*. For example, in a context in which both a pedigree-based kinship matrix and a GRM are available, it could be of interest to combine the information from both. When only a GRM is available, it could be of interest to compare application of various standard regularization approaches for covariance matrix estimation [60].

## Supplemental Data description

Supplemental Data include Supplemental Methods and 3 tables.

## Acknowledgments

This study was supported by NIH grant R01 HG001645 (to M.S.M.). The Framingham Heart Study is conducted and supported by the National Heart, Lung, and Blood Institute (NHLBI) in collaboration with Boston University (Contract No. N01-HC-25195, HHSN268201500001I and 75N92019D00031). This manuscript was not prepared in collaboration with investigators of the Framingham Heart Study and does not necessarily reflect the opinions or views of the Framingham Heart Study, Boston University, or NHLBI. Funding for SHARe Affymetrix genotyping was provided by NHLBI Contract N02-HL64278. SHARe Illumina genotyping was provided under an agreement between Illumina and Boston University. Additional funding for SABRe was provided by Division of Intramural Research, NHLBI, and Center for Population Studies, NHLBI.

## Data and code availability

- Data from the Framingham Heart Study are available at dbGAP: https://www.ncbi.nlm.nih.gov/projects/gap/cgi-bin/study.cgi?study_id=phs000007.v33.p14
- JASPER source code will be available at http://www.stat.uchicago.edu/~mcpeek/software/index.html

## Appendix A

### Proposition 1

Suppose *T* = tr(*S*_*G*_*S*_*Y*_), where 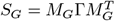, *M*_*G*_ is an *n* ×*m* matrix, Γ is an *m* ×*m* symmetric matrix, and *S*_*Y*_ is an *n* ×*n* symmetric matrix, and suppose that the following 5 conditions hold:

1. *S*_*G*_ and *S*_*Y*_ are independent (or conditionally independent given some covariates) under the null hypothesis.
2. Var_0_(vec(*M*_*G*_)) = *V*_*s*_ ⊗ *V*_*r*_, where Var_0_() represents the variance-covariance matrix under the null hypothesis, *V*_*s*_ is *m* ×*m* positive semi-definite, and *V*_*r*_ is *n* ×*n* positive semi-definite.
3. *V*_*r*_ is known (or can be estimated).
4. null space(*V*_*r*_) ⊂ [null space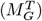 ∪ null space(*S*_*Y*_)].
5. *E*_0_(vec(*M*_*G*_)) = *µ*_*s*_ ⊗ *µ*_*r*_, where *E*_0_() represents expectation under the null hypothesis, *µ*_*s*_ is *m* ×1, *µ*_*r*_ is *n* ×*n*, and either
  a. *µ*_*s*_ = 0 or
  b. 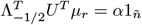 for some scalar *α* (possibly 0), where *ñ* = rank(*V*_*r*_) with *ñ* ≤ *n*, 1_*ñ*_ is the vector of length *ñ* with every entry equal to 1, *V*_*r*_ = *U* Λ*U* ^*T*^ is the eigendecomposition of *V*_*r*_, Λ_1*/*2_ is the *n* ×*ñ* matrix consisting of the *ñ* nonzero columns of the *n* ×*n* matrix Λ^1*/*2^, and Λ_−1*/*2_ is the *n* ×*ñ* matrix consisting of the *ñ* nonzero columns of the *n* ×*n* matrix (Λ^−^)^1*/*2^, where Λ^−^ is the Moore-Penrose generalized inverse of Λ (equal to Λ^−1^ when Λ is full rank).

Then the following are true:

1. We can rewrite *T* as 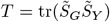, where we define the *ñ* ×*m* matrix 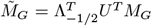 and the *ñ* ×*ñ* symmetric matrix 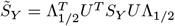, and we set 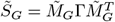.
2. Under the null hypothesis, 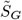 and 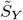are independent (or conditionally independent given the covariates).
3. Applying a permutation *σ* to the rows and columns of 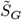 is equivalent to applying *σ* to the rows of 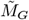 before forming 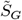, i.e., if we define 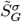 to be the matrix obtained by applying permutation *σ* to the rows and columns of 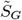, then 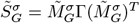, where 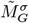 is the matrix obtained by applying permutation *σ* to the rows of 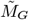.
4. Under the null hypothesis, the rows of 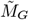 are second-order exchangeable, i.e., the following conditions hold on 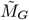:
  a. 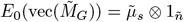, i.e., every row of 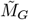 has the same mean vector 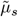 under the null hypothesis.
  b. 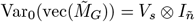, where *I*_*ñ*_ is the identity matrix of size *ñ*, i.e., every row of 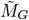 has the same covariance matrix *V*_*s*_, and the elements of different rows are uncorrelated.

**Remarks:**

1. We do not need to know or estimate *V*_*s*_.
2. If there are covariates *X* that should be conditioned on, then all the null expectations *E*_0_(·) and null variances Var_0_(·) in Proposition 1 should be taken as conditional on *X*.
3. We have assumed that both Γ and *S*_*Y*_ are symmetric. Meaningful test statistics will generally have Γ and *S*_*Y*_ both positive semi-definite as well, but that is not strictly required for our results in Propositions 1 and 2 to hold.
4. *V*_*r*_*µ*_*r*_ = 0 is sufficient for condition 5(b) to hold.
5. When *ñ* = *n*, then Λ_1*/*2_ = Λ^1*/*2^ and Λ_−1*/*2_ = Λ^−1*/*2^.
6. In practice, when an estimator 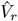 is available instead of the true matrix *V*_*r*_, we do the following:
7. Take the eigendecomposition 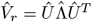, define 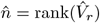, and construct 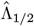 and 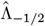 from 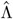 analogously to the construction of Λ_1*/*2_ and Λ_−1*/*2_ from Λ.
8. Check conditions 4 and 5 of Proposition 1 with 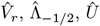, and 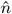 in place of *V*_*r*_, Λ_−1*/*2_, *U*, and *ñ*, respectively.
9. Define 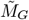and 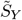 using 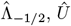, and 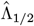 in place of Λ_−1*/*2_, *U*, and Λ_1*/*2_, respectively.

## Appendix B

From Proposition 2 below, we can obtain the first three moments of the permutation distribution of the test statistic 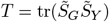 of Equation 4, given 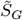 and 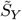, where we randomly permute the rows and columns of 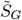, applying the same random permutation to both the rows and columns. The matrices 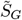and 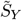 and scalar *ñ* are defined in Appendix A. To obtain the needed moments of *T*, we can set 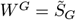 and 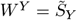 in Proposition 2.

Proposition 2 is actually a more general result that applies to a test statistic *Q*_*T*_ = tr(*W* ^*G*^*W* ^*Y*^), for any *ñ* ×*ñ* symmetric matrices *W* ^*G*^ and *W* ^*Y*^, where we permute the rows and columns of *W* ^*G*^, applying the same permutation to the rows and columns, or equivalently, we permute the rows and columns of *W* ^*Y*^, applying the same permutation to the rows and columns. (It does not matter whether we apply permutations to *W* ^*G*^ or *W* ^*Y*^ ; the permutation distribution of *Q*_*T*_ will be the same.) Proposition 2 generalizes previous results [37], which were derived for the special case when both *W* ^*G*^1_*ñ*_ = 0 and *W* ^*Y*^ 1_*ñ*_ = 0, which does not generally hold in the genetic association tests we consider when there is population structure and/or relatedness. Note that if we set 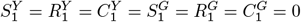 in Equation 17 and Equation 18 below, we obtain the first and second moment results of [37]. In Proposition 2, we have assumed that both *W* ^*G*^ and *W* ^*Y*^ are symmetric. Meaningful association test statistics will generally have *W* ^*G*^ and *W*_*Y*_ both positive semi-definite as well, but that is not strictly required for our results in Propositions 1 and 2 to hold.

### Proposition 2

Suppose *Q*_*T*_ = tr(*W* ^*G*^*W* ^*Y*^), where *W* ^*Y*^ and *W* ^*G*^ are both *ñ* ×*ñ* symmetric matrices. Given *W* ^*G*^ and *W* ^*Y*^, consider the distribution of *Q*_*T*_ obtained by randomly permuting the rows and columns of *W* ^*G*^, where the same permutation is applied to the rows and the columns. Let *S*_*ñ*_ denote the set of permutations of the indices (1, …, *ñ*). In the permutation distribution, we assume that each *σ* ∈ *S*_*ñ*_ has probability 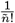. We obtain the first 3 moments of the distribution of *Q*_*T*_ given *W* ^*G*^ and *W* ^*Y*^ as follows:

1. The first moment of *Q*_*T*_ is given by

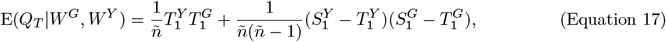

where 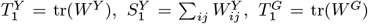, and 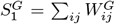, where tr(·) denotes trace, 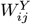 and 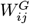 are the (*i, j*)th entries of *W* ^*Y*^ and *W* ^*G*^, respectively, and we use ∑ _*ij*_ to denote 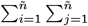.
2. The second moment of *Q*_*T*_ is given by

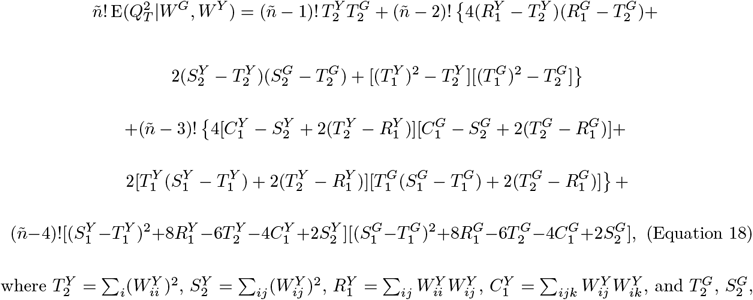

where 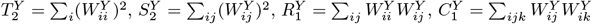, and 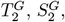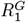 and 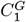 are defined analogously.
3. An expression for the third moment of *Q*_*T*_ is given in the Supplemental Methods. Like the first and second moments, the third moment is also symmetric in *W* ^*G*^ and *W* ^*Y*^ (because it does not matter whether *W* ^*G*^ or *W* ^*Y*^ is permuted), and the third moment can also can be written in terms of a small number of easily computable functions of *W* ^*Y*^ and *W* ^*G*^.

**Proof:** See Supplemental Methods.

