## Supplement for "JASPER: fast, powerful, multitrait association testing in structured samples gives insight on pleiotropy in gene expression"

### Supplemental Methods

#### Proof of Proposition 1

Proposition 1 is stated in Appendix A. Five conditions are stated, and four conclusions are reached. To prove conclusion 1, we note that

$$\tilde{M}_G^T \tilde{S}_Y \tilde{M}_G = M_G^T U \Lambda_{-1/2} \Lambda_{1/2}^T U^T S_Y U \Lambda_{1/2} \Lambda_{-1/2}^T U^T M_G,$$

where $\Lambda_{-1/2}\Lambda_{1/2}^T$ is the $n \times n$ diagonal matrix which has first $n_+$ diagonal elements equal to 1 and all other elements equal to 0, so $U\Lambda_{-1/2}\Lambda_{1/2}^T U^T = \sum_{i=1}^{n_+} u_i u_i^T$, and we get

$$\tilde{M}_G^T \tilde{S}_Y \tilde{M}_G = M_G^T \sum_{i=1}^{n_+} u_i u_i^T S_Y \sum_{i=1}^{n_+} u_i u_i^T M_G,$$

where $u_i$ is the $i$th column of $U$. Note that the null space of $V_r$ is the space spanned by the last $n - n_+$ columns of $U$. Condition (4) then implies that each $u_i$ such that $n_+ < i \le n$ must be in the null space of either $M_G$ or $S_Y$ or both, so we obtain

$$\tilde{M}_G^T \tilde{S}_Y \tilde{M}_G = M_G^T \sum_{i=1}^{n} u_i u_i^T S_Y \sum_{i=1}^{n} u_i u_i^T M_G = M_G^T S_Y M_G.$$

Conclusion 1 follows immediately from this by right-multiplying both sides by $K_G$ and using the cyclic property of trace. To prove conclusion 2, note that under condition 2, $V_s$ and $V_r$ are obtained as a result of $\mathrm{Cov}_0(\cdot)$, so they cannot be random under the null distribution, so the conclusion that $\tilde{S}_G$ and $\tilde{S}_Y$ are independent (or conditionally independent given some covariates) under the null hypothesis follows trivially from condition 1. To verify conclusion 3, let $P_\sigma$ denote the orthogonal $\tilde{n} \times \tilde{n}$ matrix such that for all $k \ge 1$ and all $\tilde{n} \times k$ matrices $A$, $P_\sigma A$ is the matrix $A$ with permutation $\sigma$ applied to its rows. Then we have $\tilde{S}_G^\sigma = P_\sigma \tilde{S}_G P_\sigma^T$ which, by the associative property of matrix multiplication, is equal to $\tilde{M}_G^\sigma K_G (\tilde{M}_G^\sigma)^T$, where $\tilde{M}_G^\sigma = P_\sigma \tilde{M}_G$. Next we verify conclusion 4(a). Note that from a basic property of the vec operator, we have

$$\mathrm{vec}(\tilde{M}_G) = (I_m \otimes (\Lambda_{-1/2}^T U^T))\mathrm{vec}(M_G), \qquad \text{(Equation S1)}$$

so condition 5 implies

$$E_0(\mathrm{vec}(\tilde{M}_G)) = (I_m \otimes (\Lambda_{-1/2}^T U^T))(\mu_s \otimes \mu_r) = \mu_s \otimes (\Lambda_{-1/2}^T U^T \mu_r),$$

where either $\mu_s = 0$ or $\Lambda_{-1/2}^T U^T \mu_r = \alpha 1_{n+}$. If $\mu_s = 0$, then we get conclusion 4(a) with $\tilde{\mu}_s = 0$. If $\Lambda_{-1/2}^T U^T \mu_r = \alpha 1_{n+}$, then we get conclusion 4(a) with $\tilde{\mu}_s = \alpha \mu_s$. Next we verify conclusion 4(b). From condition 2 and Equation S1, we get

$$\mathrm{Var}_0(\mathrm{vec}(\tilde{M}_G)) = V_s \otimes (\Lambda_{-1/2}^T U^T V_r U \Lambda_{-1/2}) = V_s \otimes (\Lambda_{-1/2}^T \Lambda \Lambda_{-1/2}) = V_s \otimes I_{n_+},$$

which proves conclusion 4(b). □

#### Proposition 2 Part 3

Proposition 2 is stated in Appendix B, except for part 3, which is as follows:

3. The third moment of $Q_T$ is given by

$$\begin{aligned}
&\tilde{n}!\,\mathrm{E}(Q_T^3 | W^G, W^Y) \\
&= (\tilde{n}-1)! \sum_i (W_{ii}^Y)^3 \sum_i (W_{ii}^G)^3 \\
&+ (\tilde{n}-2)! \Bigg\{ 6 \sum_{ij}{}' W_{ii}^Y W_{jj}^Y W_{ij}^Y \cdot \sum_{ij}{}' W_{ii}^G W_{jj}^G W_{ij}^G + 12 \sum_{ij}{}' W_{ii}^Y (W_{ij}^Y)^2 \cdot \\
&\sum_{ij}{}' W_{ii}^G (W_{ij}^G)^2 + 6 \sum_{ij}{}' (W_{ii}^Y)^2 W_{ij}^Y \cdot \sum_{ij}{}' (W_{ii}^G)^2 W_{ij}^G + 3 \sum_{ij}{}' (W_{ii}^Y)^2 W_{jj}^Y \cdot \\
&\sum_{ij}{}' (W_{ii}^G)^2 W_{jj}^G + 4 \sum_{ij}{}' (W_{ij}^Y)^3 \cdot \sum_{ij}{}' (W_{ij}^G)^3 \Bigg\} \\
&+ (\tilde{n}-3)! \Bigg\{ \sum_{ijk}{}' W_{ii}^Y W_{jj}^Y W_{kk}^Y \cdot \sum_{ijk}{}' W_{ii}^G W_{jj}^G W_{kk}^G + 12 \sum_{ijk}{}' W_{ii}^Y W_{jj}^Y W_{ik}^Y \cdot \\
&\sum_{ijk}{}' W_{ii}^G W_{jj}^G W_{ik}^G + 12 \sum_{ijk}{}' W_{ii}^Y W_{ij}^Y W_{ik}^Y \cdot \sum_{ijk}{}' W_{ii}^G W_{ij}^G W_{ik}^G + 24 \sum_{ijk}{}' W_{ii}^Y W_{ij}^Y W_{jk}^Y \cdot \\
&\sum_{ijk}{}' W_{ii}^G W_{ij}^G W_{jk}^G + 8 \sum_{ijk}{}' W_{ij}^Y W_{ik}^Y W_{jk}^Y \cdot \sum_{ijk}{}' W_{ij}^G W_{ik}^G W_{jk}^G + 6 \sum_{ijk}{}' W_{ii}^Y (W_{jk}^Y)^2 \cdot \\
&\sum_{ijk}{}' W_{ii}^G (W_{jk}^G)^2 + 3 \sum_{ijk}{}' (W_{ii}^Y)^2 W_{jk}^Y \cdot \sum_{ijk}{}' (W_{ii}^G)^2 W_{jk}^G + \\
&24 \sum_{ijk}{}' (W_{ij}^Y)^2 W_{ik}^Y \cdot \sum_{ijk}{}' (W_{ij}^G)^2 W_{ik}^G \Bigg\}
\end{aligned}$$

$$
\begin{aligned}
&+ (\tilde{n}-4)! \Bigg\{ 3 \sum_{ijkl}{}' W^Y_{ii} W^Y_{jj} W^Y_{kl} \cdot \sum_{ijkl}{}' W^G_{ii} W^G_{jj} W^G_{kl} + 12 \sum_{ijkl}{}' W^Y_{ii} W^Y_{ij} W^Y_{kl} \cdot \\
&\sum_{ijkl}{}' W^G_{ii} W^G_{ij} W^G_{kl} + 12 \sum_{ijkl}{}' W^Y_{ii} W^Y_{jk} W^Y_{jl} \cdot \sum_{ijkl}{}' W^G_{ii} W^G_{jk} W^G_{jl} + 8 \sum_{ijkl}{}' W^Y_{ij} W^Y_{ik} W^Y_{il} \cdot \\
&\sum_{ijkl}{}' W^G_{ij} W^G_{ik} W^G_{il} + 24 \sum_{ijkl}{}' W^Y_{ij} W^Y_{ik} W^Y_{jl} \cdot \sum_{ijkl}{}' W^G_{ij} W^G_{ik} W^G_{jl} + \\
&6 \sum_{ijkl}{}' (W^Y_{ij})^2 W^Y_{kl} \cdot \sum_{ijkl}{}' (W^G_{ij})^2 W^G_{kl} \Bigg\} \\
&+ (\tilde{n}-5)! \Bigg\{ 12 \sum_{ijklm}{}' W^Y_{ij} W^Y_{ik} W^Y_{lm} \cdot \sum_{ijklm}{}' W^G_{ij} W^G_{ik} W^G_{lm} + \\
&3 \sum_{ijklm}{}' W^Y_{ii} W^Y_{jk} W^Y_{lm} \cdot \sum_{ijklm}{}' W^G_{ii} W^G_{jk} W^G_{lm} \Bigg\} \\
&+ (\tilde{n}-6)! \sum_{ijklmo}{}' W^Y_{ij} W^Y_{kl} W^Y_{mo} \cdot \sum_{ijklmo}{}' W^G_{ij} W^G_{kl} W^G_{mo}, \qquad \text{(Equation S2)}
\end{aligned}
$$

where all summations are from 1 to $\tilde{n}$, $\sum'$ denotes summation over distinct indices, e.g.,

$$
\sum_{i,j}{}' \text{ denotes } \sum_i \sum_{j \neq i} \text{ and } \sum_{i,j,k}{}' \text{ denotes } \sum_i \sum_{j \neq i} \sum_{\substack{k \neq i \\ k \neq j}}, \text{ etc.,}
$$

and where the expression in Equation S2 can be computed in terms of a small number of easily obtainable matrix quantities (see Lemma 4).

#### Proof of Proposition 2

1. To prove part 1 of Proposition 2, we first define $W^{G,\sigma}$ to be the matrix obtained by applying permutation $\sigma \in S_{\tilde{n}}$ to both the rows and columns of $W^G$. Then we have

$$
\begin{aligned}
\mathrm{E}(Q_T | W^G, W^Y) &= \frac{1}{\tilde{n}!} \sum_{\sigma \in S_{\tilde{n}}} \mathrm{tr}(W^{G,\sigma} W^Y) \\
&= \frac{1}{\tilde{n}!} \sum_{ij} \sum_{\sigma \in S_{\tilde{n}}} W^{G,\sigma}_{ij} W^Y_{ij} \\
&= \frac{1}{\tilde{n}!} \sum_i W^Y_{ii} \sum_{\sigma \in S_{\tilde{n}}} W^{G,\sigma}_{ii} + \frac{1}{\tilde{n}!} \sum_{i \neq j} W^Y_{ij} \sum_{\sigma \in S_{\tilde{n}}} W^{G,\sigma}_{ij}.
\end{aligned}
$$

Part 1 of Proposition 2 then follows from Lemma 1 below. (Note that the proof of part 1 does not require symmetry of either $W^G$ or $W^Y$.) □

**Lemma 1.** *For any $\tilde{n} \times \tilde{n}$ matrix $A$ (not necessarily symmetric) and any $\sigma \in S_{\tilde{n}}$, let $A^\sigma$ denote the matrix obtained by applying permutation $\sigma$ to both the rows and columns of $A$. Then for up to two distinct indices $i$ and $j$, we have*

$$\sum_{\sigma \in S_{\tilde{n}}} A^\sigma_{ii} = (\tilde{n}-1)! \sum_i A_{ii} = (\tilde{n}-1)! \mathrm{tr}(A),$$

$$\sum_{\sigma \in S_{\tilde{n}}} A^\sigma_{ij} = (\tilde{n}-2)! \sum_{i \neq j} A_{ij} = (\tilde{n}-2)! \left[ \sum_{ij} A_{ij} - \mathrm{tr}(A) \right].$$

2. To prove part 2 of Proposition 2, note that

$$\begin{aligned}
&\tilde{n}!\, \mathrm{E}(Q_T^2 | W^G, W^Y) \\
&= \sum_{\sigma \in S_{\tilde{n}}} \mathrm{tr}(W^{G,\sigma} W^Y)^2 \\
&= \sum_{ijkl} \sum_{\sigma \in S_{\tilde{n}}} W^{G,\sigma}_{ij} W^Y_{ij} W^{G,\sigma}_{kl} W^Y_{kl} \\
&= \sum_{i} \sum_{\sigma \in S_{\tilde{n}}} (W^Y_{ii})^2 (W^{G,\sigma}_{ii})^2 \\
&+ \sum_{ij}{}' \sum_{\sigma \in S_{\tilde{n}}} \left\{ 4 W^Y_{ii} W^Y_{ij} W^{G,\sigma}_{ii} W^{G,\sigma}_{ij} + 2 (W^Y_{ij})^2 (W^{G,\sigma}_{ij})^2 + W^Y_{ii} W^Y_{jj} W^{G,\sigma}_{ii} W^{G,\sigma}_{jj} \right\} \\
&+ \sum_{ijk}{}' \sum_{\sigma \in S_{\tilde{n}}} \left\{ 4 W^Y_{ij} W^Y_{ik} W^{G,\sigma}_{ij} W^{G,\sigma}_{ik} + 2\, W^Y_{ii} W^Y_{jk} W^{G,\sigma}_{ii} W^{G,\sigma}_{jk} \right\} \\
&+ \sum_{ijkl}{}' \sum_{\sigma \in S_{\tilde{n}}} W^Y_{ij} W^Y_{kl} W^{G,\sigma}_{ij} W^{G,\sigma}_{kl}, \qquad \text{(Equation S3)}
\end{aligned}$$

where this uses symmetry of both $W^G$ and $W^Y$. Using Lemma 2 below, we obtain

$$\begin{aligned}
&\tilde{n}!\, \mathrm{E}(Q_T^2 | W^G W^Y) \\
&= (\tilde{n}-1)! \sum_i (W^Y_{ii})^2 \cdot \sum_i (W^G_{ii})^2 \\
&+ (\tilde{n}-2)! \left\{ 4 \sum_{ij}{}' W^Y_{ii} W^Y_{ij} \cdot \sum_{ij}{}' W^G_{ii} W^G_{ij} + 2 \sum_{ij}{}' (W^Y_{ij})^2 \cdot \sum_{ij}{}' (W^G_{ij})^2 + \right. \\
&\left. \sum_{ij}{}' W^Y_{ii} W^Y_{jj} \cdot \sum_{ij}{}' W^G_{ii} W^G_{jj} \right\} \\
&+ (\tilde{n}-3)! \left\{ 4 \sum_{ijk}{}' W^Y_{ij} W^Y_{ik} \cdot \sum_{ijk}{}' W^G_{ij} W^G_{ik} + 2 \sum_{ijk}{}' W^Y_{ii} W^Y_{jk} \cdot \sum_{ijk}{}' W^G_{ii} W^G_{jk} \right\}
\end{aligned}$$

$$+ (\tilde{n}-4)! {\sum_{ijkl}}' W^Y_{ij} W^Y_{kl} \cdot {\sum_{ijkl}}' W^G_{ij} W^G_{kl}, \qquad \text{(Equation S4)}$$

and part 2 of Proposition 2 follows using Lemma 4 below. □

**Lemma 2.** *In the same setting as Lemma 1, for up to four distinct indices $i, j, k$ and $l$, we have*

$$\sum_{\sigma \in S_{\tilde{n}}} (A^\sigma_{ii})^2 = (\tilde{n}-1)! \sum_i A^2_{ii}, \quad \sum_{\sigma \in S_{\tilde{n}}} A^\sigma_{ii} A^\sigma_{ij} = (\tilde{n}-2)! {\sum_{ij}}' A_{ii} A_{ij},$$
$$\sum_{\sigma \in S_{\tilde{n}}} (A^\sigma_{ij})^2 = (\tilde{n}-2)! {\sum_{ij}}' A^2_{ij}, \quad \sum_{\sigma \in S_{\tilde{n}}} A^\sigma_{ii} A^\sigma_{jj} = (\tilde{n}-2)! {\sum_{ij}}' A_{ii} A_{jj},$$
$$\sum_{\sigma \in S_{\tilde{n}}} A^\sigma_{ij} A^\sigma_{ik} = (\tilde{n}-3)! {\sum_{ijk}}' A_{ij} A_{ik}, \quad \sum_{\sigma \in S_{\tilde{n}}} A^\sigma_{ii} A^\sigma_{jk} = (\tilde{n}-3)! {\sum_{ijk}}' A_{ii} A_{jk},$$
$$\sum_{\sigma \in S_{\tilde{n}}} A^\sigma_{ij} A^\sigma_{kl} = (\tilde{n}-4)! {\sum_{ijkl}}' A_{ij} A_{kl}.$$

3. To prove part 3 of Proposition 2, we note that

$$\tilde{n}!\, \mathrm{E}(Q^3_T | W^G, W^Y)$$
$$= \sum_{\sigma \in S_{\tilde{n}}} \mathrm{tr}(W^{G,\sigma} W^Y)^3$$
$$= \sum_{ijklmo} \sum_{\sigma \in S_{\tilde{n}}} W^{G,\sigma}_{ij} W^Y_{ij} W^{G,\sigma}_{kl} W^Y_{kl} W^{G,\sigma}_{mo} W^Y_{mo}$$
$$= \sum_i \sum_{\sigma \in S_{\tilde{n}}} (W^Y_{ii})^3 (W^{G,\sigma}_{ii})^3$$
$$+ {\sum_{ij}}' \sum_{\sigma \in S_{\tilde{n}}} \Big\{ 6 W^Y_{ii} W^Y_{jj} W^Y_{ij} W^{G,\sigma}_{ii} W^{G,\sigma}_{jj} W^{G,\sigma}_{ij} + 12 W^Y_{ii} (W^Y_{ij})^2 W^{G,\sigma}_{ii} (W^{G,\sigma}_{ij})^2 +$$
$$6 (W^Y_{ii})^2 W^Y_{ij} (W^{G,\sigma}_{ii})^2 W^{G,\sigma}_{ij} + 3 (W^Y_{ii})^2 W^Y_{jj} (W^{G,\sigma}_{ij})^2 W^{G,\sigma}_{jj} + 4 (W^Y_{ij})^3 (W^{G,\sigma}_{ij})^3 \Big\}$$
$$+ {\sum_{ijk}}' \sum_{\sigma \in S_{\tilde{n}}} \Big\{ W^Y_{ii} W^Y_{jj} W^Y_{kk} W^{G,\sigma}_{ii} W^{G,\sigma}_{jj} W^{G,\sigma}_{kk} + 12 W^Y_{ii} W^Y_{jj} W^Y_{ik} W^{G,\sigma}_{ii} W^{G,\sigma}_{jj} W^{G,\sigma}_{ik} +$$
$$12 W^Y_{ii} W^Y_{ij} W^Y_{ik} W^{G,\sigma}_{ii} W^{G,\sigma}_{ij} W^{G,\sigma}_{ik} + 24 W^Y_{ii} W^Y_{ij} W^Y_{jk} W^{G,\sigma}_{ii} W^{G,\sigma}_{ij} W^{G,\sigma}_{jk} +$$
$$8 W^Y_{ij} W^Y_{ik} W^Y_{jk} W^{G,\sigma}_{ij} W^{G,\sigma}_{ik} W^{G,\sigma}_{jk} + 6 W^Y_{ii} (W^Y_{jk})^2 W^{G,\sigma}_{ii} (W^{G,\sigma}_{jk})^2 +$$
$$3 (W^Y_{ii})^2 W^Y_{jk} (W^{G,\sigma}_{ii})^2 W^{G,\sigma}_{jk} + 24 (W^Y_{ij})^2 W^Y_{ik} (W^{G,\sigma}_{ij})^2 W^{G,\sigma}_{ik} \Big\}$$
$$+ {\sum_{ijkl}}' \sum_{\sigma \in S_{\tilde{n}}} \Big\{ 3 W^Y_{ii} W^Y_{jj} W^Y_{kl} W^{G,\sigma}_{ii} W^{G,\sigma}_{jj} W^{G,\sigma}_{kl} + 12 W^Y_{ii} W^Y_{ij} W^Y_{kl} W^{G,\sigma}_{ii} W^{G,\sigma}_{ij} W^{G,\sigma}_{kl} +$$
$$12 W^Y_{ii} W^Y_{jk} W^Y_{jl} W^{G,\sigma}_{ii} W^{G,\sigma}_{jk} W^{G,\sigma}_{jl} + 8 W^Y_{ij} W^Y_{ik} W^Y_{il} W^{G,\sigma}_{ij} W^{G,\sigma}_{ik} W^{G,\sigma}_{il} +$$

$$24W^Y_{ij}W^Y_{ik}W^Y_{jl}W^{G,\sigma}_{ij}W^{G,\sigma}_{ik}W^{G,\sigma}_{jl} + 6(W^Y_{ij})^2W^Y_{kl}(W^{G,\sigma}_{ij})^2W^{G,\sigma}_{kl}\Big\}$$

$$+ \sum_{ijklm}{}' \sum_{\sigma\in S_{\tilde n}} \Big\{12W^Y_{ij}W^Y_{ik}W^Y_{lm}W^{G,\sigma}_{ij}W^{G,\sigma}_{ik}W^{G,\sigma}_{lm} + 3W^Y_{ii}W^Y_{jk}W^Y_{lm}W^{G,\sigma}_{ii}W^{G,\sigma}_{jk}W^{G,\sigma}_{lm}\Big\}$$

$$+ \sum_{ijklmo}{}' \sum_{\sigma\in S_{\tilde n}} W^Y_{ij}W^Y_{kl}W^Y_{mo}W^{G,\sigma}_{ij}W^{G,\sigma}_{kl}W^{G,\sigma}_{mo}. \qquad \text{(Equation S5)}$$

Using Lemma 3, the desired result follows. □

**Lemma 3.** *In the same setting as Lemma 1, for up to six distinct indices $i, j, k, l, m$ and $o$, we have*

$$\sum_{\sigma\in S_{\tilde n}} (A^\sigma_{ii})^3 = (\tilde n - 1)! \sum_i A^3_{ii}, \quad \sum_{\sigma\in S_{\tilde n}} A^\sigma_{ii}A^\sigma_{jj}A^\sigma_{ij} = (\tilde n - 2)! \sum_{ij}{}' A_{ii}A_{jj}A_{ij},$$

$$\sum_{\sigma\in S_{\tilde n}} A^\sigma_{ii}(A^\sigma_{ij})^2 = (\tilde n - 2)! \sum_{ij}{}' A_{ii}A^2_{ij}, \quad \sum_{\sigma\in S_{\tilde n}} (A^\sigma_{ii})^2A^\sigma_{ij} = (\tilde n - 2)! \sum_{ij}{}' A^2_{ii}A_{ij},$$

$$\sum_{\sigma\in S_{\tilde n}} (A^\sigma_{ii})^2A^\sigma_{jj} = (\tilde n - 2)! \sum_{ij}{}' A^2_{ii}A_{jj}, \quad \sum_{\sigma\in S_{\tilde n}} (A^\sigma_{ij})^3 = (\tilde n - 2)! \sum_i A^3_{ij},$$

$$\sum_{\sigma\in S_{\tilde n}} A^\sigma_{ii}A^\sigma_{jj}A^\sigma_{kk} = (\tilde n - 3)! \sum_{ijk}{}' A_{ii}A_{jj}A_{kk}, \quad \sum_{\sigma\in S_{\tilde n}} A^\sigma_{ii}A^\sigma_{jj}A^\sigma_{ik} = (\tilde n - 3)! \sum_{ijk}{}' A_{ii}A_{jj}A_{ik},$$

$$\sum_{\sigma\in S_{\tilde n}} A^\sigma_{ii}A^\sigma_{ij}A^\sigma_{ik} = (\tilde n - 3)! \sum_{ijk}{}' A_{ii}A_{ij}A_{ik}, \quad \sum_{\sigma\in S_{\tilde n}} A^\sigma_{ii}A^\sigma_{ij}A^\sigma_{jk} = (\tilde n - 3)! \sum_{ijk}{}' A_{ii}A_{ij}A_{jk},$$

$$\sum_{\sigma\in S_{\tilde n}} A^\sigma_{ij}A^\sigma_{ik}A^\sigma_{jk} = (\tilde n - 3)! \sum_{ijk}{}' A_{ij}A_{ik}A_{jk}, \quad \sum_{\sigma\in S_{\tilde n}} A^\sigma_{ii}(A^\sigma_{jk})^2 = (\tilde n - 3)! \sum_{ijk}{}' A_{ii}A^2_{jk},$$

$$\sum_{\sigma\in S_{\tilde n}} (A^\sigma_{ii})^2A^\sigma_{jk} = (\tilde n - 3)! \sum_{ijk}{}' A^2_{ii}A_{jk}, \quad \sum_{\sigma\in S_{\tilde n}} (A^\sigma_{ij})^2A^\sigma_{ik} = (\tilde n - 3)! \sum_{ijk}{}' A^2_{ij}A_{ik},$$

$$\sum_{\sigma\in S_{\tilde n}} A^\sigma_{ii}A^\sigma_{jj}A^\sigma_{kl} = (\tilde n - 4)! \sum_{ijkl}{}' A_{ii}A_{jj}A_{kl}, \quad \sum_{\sigma\in S_{\tilde n}} A^\sigma_{ii}A^\sigma_{ij}A^\sigma_{kl} = (\tilde n - 4)! \sum_{ijkl}{}' A_{ii}A_{ij}A_{kl},$$

$$\sum_{\sigma\in S_{\tilde n}} A^\sigma_{ii}A^\sigma_{jk}A^\sigma_{jl} = (\tilde n - 4)! \sum_{ijkl}{}' A_{ii}A_{jk}A_{jl}, \quad \sum_{\sigma\in S_{\tilde n}} A^\sigma_{ij}A^\sigma_{ik}A^\sigma_{il} = (\tilde n - 4)! \sum_{ijkl}{}' A_{ij}A_{ik}A_{il},$$

$$\sum_{\sigma\in S_{\tilde n}} A^\sigma_{ij}A^\sigma_{ik}A^\sigma_{jl} = (\tilde n - 4)! \sum_{ijkl}{}' A_{ij}A_{ik}A_{jl}, \quad \sum_{\sigma\in S_{\tilde n}} (A^\sigma_{ij})^2A^\sigma_{kl} = (\tilde n - 4)! \sum_{ijkl}{}' A^2_{ij}A_{kl},$$

$$\sum_{\sigma\in S_{\tilde n}} A^\sigma_{ij}A^\sigma_{ik}A^\sigma_{lm} = (\tilde n - 5)! \sum_{ijklm}{}' A_{ij}A_{ik}A_{lm}, \quad \sum_{\sigma\in S_{\tilde n}} A^\sigma_{ii}A^\sigma_{jk}A^\sigma_{lm} = (\tilde n - 5)! \sum_{ijklm}{}' A_{ii}A_{jk}A_{lm},$$

$$\sum_{\sigma\in S_{\tilde n}} A^\sigma_{ik}A^\sigma_{kl}A^\sigma_{mo} = (\tilde n - 6)! \sum_{ijklmo}{}' A_{ij}A_{kl}A_{mo}.$$

**Lemma 4.** *All the sums needed in Proposition 2 can be expressed in terms of the following* *quantities,*

$$T_1^A = \sum_i A_{ii},\ T_2^A = \sum_i A_{ii}^2,\ T_3^A = \sum_i A_{ii}^3,$$

$$S_1^A = \sum_{ij} A_{ij},\ S_2^A = \sum_{ij} A_{ij}^2,\ S_3^A = \sum_{ij} A_{ij}^3,$$

$$R_1^A = \sum_{ij} A_{ii}A_{ij},\ R_2^A = \sum_{ij} A_{ii}A_{ij}^2,\ R_3^A = \sum_{ij} A_{ii}^2 A_{ij},\ R_4^A = \sum_{ij} A_{ii}A_{jj}A_{ij},$$

$$C_1^A = \sum_{ijk} A_{ij}A_{ik},\ C_2^A = \sum_{ijk} A_{ij}^2 A_{ik},\ C_3^A = \sum_{ijk} A_{ii}A_{ij}A_{ik},\ C_4^A = \sum_{ijk} A_{ii}A_{ij}A_{jk},$$

$$C_5^A = \sum_{ijk} A_{ij}A_{ik}A_{jk},\ D_1^A = \sum_{ijkl} A_{ij}A_{ik}A_{il},\ D_2^A = \sum_{ijkl} A_{ij}A_{ik}A_{jl},$$

*where the matrix $A$ can be replaced by $W^G$ or $W^Y$.*

*Proof.*

$$\sum_{ij}{}' A_{ij} = S_1^A - T_1^A,$$

$$\sum_{ij}{}' A_{ij}^2 = S_2^A - T_2^A,$$

$$\sum_{ij}{}' A_{ij}^3 = S_3^A - T_3^A,$$

$$\sum_{ij}{}' A_{ii}A_{ij} = R_1^A - T_2^A,$$

$$\sum_{ij}{}' A_{ii}A_{ij}^2 = R_2^A - T_3^A,$$

$$\sum_{ij}{}' A_{ii}^2 A_{ij} = R_3^A - T_3^A,$$

$$\sum_{ij}{}' A_{ii}A_{jj} = (T_1^A)^2 - T_2^A,$$

$$\sum_{ij}{}' A_{ii}^2 A_{jj} = T_1^A T_2^A - T_3^A,$$

$$\sum_{ij}{}' A_{ii}A_{jj}A_{ij} = R_4^A - T_3^A,$$

$$\sum_{ijk}{}' A_{ii}A_{jk} = T_1^A(S_1^A - T_1^A) + 2(T_2^A - R_1^A),$$

$$\sum_{ijk}{}' A_{ii}A_{jk}^2 = T_1^A(S_2^A - T_2^A) + 2(T_3^A - R_2^A),$$

$$\sum_{ijk}{}' A_{ii}^2 A_{jk} = T_2^A(S_1^A - T_1^A) + 2(T_3^A - R_3^A),$$

$$\sum_{ijk}{}' A_{ij} A_{ik} = C_1^A - S_2^A + 2(T_2^A - R_1^A),$$

$$\sum_{ijk}{}' A_{ij}^2 A_{ik} = C_2^A - R_2^A - R_3^A - S_3^A + 2T_3^A,$$

$$\sum_{ijk}{}' A_{ii} A_{ij} A_{jk} = C_4^A - R_2^A - R_3^A - R_4^A + 2T_3^A,$$

$$\sum_{ijk}{}' A_{ii} A_{ij} A_{ik} = C_3^A - R_2^A + 2(T_3^A - R_3^A),$$

$$\sum_{ijk}{}' A_{ii} A_{jj} A_{ik} = T_1^A(R_1^A - T_2^A) - R_3^A - R_4^A + 2T_3^A,$$

$$\sum_{ijk}{}' A_{ii} A_{jj} A_{kk} = T_1^A[(T_1^A)^2 - T_2^A] + 2(T_3^A - T_1^A T_2^A),$$

$$\sum_{ijk}{}' A_{ij} A_{ik} A_{jk} = C_5^A - R_2^A + 2(T_3^A - R_2^A),$$

$$\sum_{ijkl}{}' A_{ij} A_{kl} = S_1^A(S_1^A - T_1^A) + 2(R_1^A - C_1^A) - 2\sum_{ijk}{}' A_{ij} A_{ik} - \sum_{ijk}{}' A_{ii} A_{jk},$$

$$\sum_{ijkl}{}' A_{ij}^2 A_{kl} = S_2^A(S_1^A - T_1^A) + 2(R_2^A - C_2^A) - 2\sum_{ijk}{}' A_{ij}^2 A_{ik} - \sum_{ijk}{}' A_{ii}^2 A_{jk},$$

$$\sum_{ijkl}{}' A_{ii} A_{jj} A_{kl} = T_1^A \sum_{ijk}{}' A_{ii} A_{jk} - 2\sum_{ijk}{}' A_{ii} A_{jj} A_{ik} - \sum_{ijk}{}' A_{ii}^2 A_{jk},$$

$$\sum_{ijkl}{}' A_{ii} A_{ij} A_{kl} = R_1^A(S_1^A - T_1^A) + 2(R_3^A - C_3^A) - 2\sum_{ijk}{}' A_{ii} A_{ij} A_{jk} - \sum_{ijk}{}' A_{ii}^2 A_{jk},$$

$$\sum_{ijkl}{}' A_{ii} A_{jk} A_{jl} = T_1^A \sum_{ijk}{}' A_{ij} A_{ik} - 2\sum_{ijk}{}' A_{ii} A_{ij} A_{jk} - \sum_{ijk}{}' A_{ii} A_{ij} A_{ik},$$

$$\sum_{ijkl}{}' A_{ij} A_{ik} A_{il} = D_1^A - 3C_2^A + 2S_3^A - 3\sum_{ijk}{}' A_{ii} A_{ij} A_{ik},$$

$$\begin{aligned}\sum_{ijkl}{}' A_{ij} A_{ik} A_{jl} &= D_2^A - C_2^A - C_4^A - C_5^A + 2R_2^A - \sum_{ijk}{}' A_{ii} A_{ij} A_{ik} \\ &- \sum_{ijk}{}' A_{ii} A_{ij} A_{jk} - \sum_{ijk}{}' A_{ij}^2 A_{ik},\end{aligned}$$

$$\sum_{ijklm}{}' A_{ii} A_{jk} A_{lm} = T_1^A \sum_{ijkl}{}' A_{ij} A_{kl} - 4\sum_{ijkl}{}' A_{ii} A_{ij} A_{kl},$$

$$\begin{aligned}\sum_{ijklm}{}' A_{ij} A_{ik} A_{lm} &= (C_1^A - S_2^A)(S_1^A - T_1^A) + 4(C_2^A + C_4^A - D_2^A) + 2C_5^A - 6R_2^A \\ &- 2\sum_{ijkl}{}' A_{ii} A_{ij} A_{kl} - 2\sum_{ijkl}{}' A_{ij} A_{ik} A_{il},\end{aligned}$$

$$\sum_{ijklmo}{}' A_{ij}A_{kl}A_{mo} = S_1^A \sum_{ijkl}{}' A_{ij}A_{kl} + 4(R_1^A - C_1^A)(S_1^A - T_1^A) + 8(D_1^A + D_2^A - C_2^A - C_4^A) + 24(R_3^A - C_3^A) - 4\sum_{ijklm}{}' A_{ij}A_{ik}A_{lm} - \sum_{ijklm}{}' A_{ii}A_{jk}A_{lm}.$$

□

#### Proof that the score test for association based on Equation 12 is of the form in Equation 1

In the main text, we state that the score test for the null hypothesis $H_0 : \tau^2 = 0$ in the model in Equation 12 can be written in the form of Equation 1 where $S_G$ is as given in Equation 10 and $S_Y = \tilde{Y}\hat{V}_g\tilde{Y}^T$, where $\tilde{Y}$ is as given in Equation 13. (These all refer to equations in the main text.)

*Proof.* Let $l$ denote the log-likelihood for the model of Equation 12 and note that

$$\left.\frac{\partial l}{\partial \tau^2}\right|_{\tau^2=0} = -\frac{1}{2}\mathrm{tr}(\Omega_0^{-1}[V_g \otimes (\tilde{G}W\tilde{G}^T)]) + \frac{1}{2}\mathrm{vec}(Y-\mu)^T\Omega_0^{-1}[V_g \otimes (\tilde{G}W\tilde{G}^T)]\Omega_0^{-1}\mathrm{vec}(Y-\mu),$$

where $\Omega_0 = V_a \otimes K + V_e \otimes I_n$. Using the common convention, we neglect the first term, which subtracts off the null mean of the test statistic, because it does not involve $Y$ so would have no impact on the JASPER assessment of significance. Thus, plugging in the estimated values of the remaining parameters under the null hypothesis, using the definition of $\tilde{Y}$ given in Equation 13 of the main text, and neglecting the factor of 1/2, we can take the score statistic to be

$$T = \mathrm{vec}(\tilde{Y})^T[\hat{V}_g \otimes (\tilde{G}W\tilde{G}^T)]\mathrm{vec}(\tilde{Y}).$$

We use the identity $\mathrm{vec}(ABC) = (C^T \otimes A)\mathrm{vec}(B)$ to get

$$[\hat{V}_g \otimes (\tilde{G}W\tilde{G}^T)]\mathrm{vec}(\tilde{Y}) = \mathrm{vec}[(\tilde{G}W\tilde{G}^T)\tilde{Y}\hat{V}_g],$$

from which we get

$$T = \mathrm{vec}(\tilde{Y})^T\mathrm{vec}[(\tilde{G}W\tilde{G}^T)\tilde{Y}\hat{V}_g]. \qquad \text{(Equation S6)}$$

Then apply the identity $\text{tr}(A^T B) = \text{vec}(A)^T \text{vec}(B)$ to Equation S6 to obtain

$$\begin{aligned} T &= \text{tr}[\tilde{Y}^T(\tilde{G}W\tilde{G}^T)\tilde{Y}\hat{V}_g] \\ &= \text{tr}[(\tilde{G}W\tilde{G}^T)\tilde{Y}\hat{V}_g\tilde{Y}^T] \\ &= \text{tr}(S_G S_Y), \end{aligned}$$

where the second equality is by the cyclic property of trace. □

#### Further Details on Covariates and Trait Models for Simulations

Three observed covariates are simulated: sex, age and a continuous covariate with the standard Gaussian distribution. When no familial relatedness is present in the sample, sex is generated as a Bernoulli(0.5) variable and age is a variable uniformly distributed between 20 and 60. When the sample consists of related individuals, sex is determined by the pedigree structure shown in Figure S1. In addition, age is a variable uniformly distributed within 1.5 years of 73, 75, 46, 43, 40, 46, 40, 43, 47, 51, 18, 21, 15, 15, 12, 9, 13, 17, 24, 21, 18, and 14 years respectively for individuals labeled $1 - 22$ in a given pedigree whose structure is shown in Figure S1. For all simulation scenarios, the covariate values are assumed to be independent across individuals and are regenerated for each replicate.

**Figure S1:** Three-generation pedigree of 22 individuals used in the simulation studies.

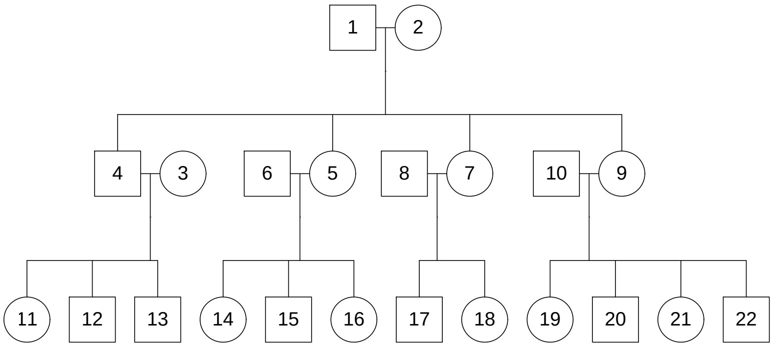


For Model I of Equation 15, the parameters $\beta, \delta_1, \delta_2, \zeta, \sigma_a^2$ and $\sigma_e^2$ are chosen to satisfy the following conditions for each trait: (1) The total trait variance is set to 1 and the mean to 0; (2) The three non-trivial covariates in $X$ each explain an equal amount of the total trait variance; (3) The unobserved major genes each explain 1.5% of the total trait variance; (4) The total variance explained by the ancestry effect ($a\zeta$) and additive polygenic effect ($\alpha$) is set to 20% of the total trait variance; (5) For the variance in (4), either it is completely explained by the ancestry fixed effect (in the case of no relatedness), or half of it is explained by the ancestry effect with the remaining half explained by the polygenic additive effect (in the case of relatedness) (6) The random error $\epsilon$ explains 20% of the total trait variance; (7) Given the variance explained by $G\gamma$, the details of which are explained in the next two paragraphs, covariates explain the remaining amount of trait variance. For Model II in Equation 16, we modify condition (6) so that the random error $\epsilon$ explains 60% of the total trait variance (on the liability scale), and keep the other conditions the same but applied on the liability scale.

For each given setting, the 100,000 variants used to correct for population structure are re-simulated three times and in each simulation, the GRM estimate $K$ is given by

$$K_{ij} = \frac{1}{L}\sum_{l=1}^{L}\frac{(G_{il}-2\hat{p}_l)(G_{jl}-2\hat{p}_l)}{2\hat{p}_l(1-\hat{p}_l)}, \text{ where } L = 10^5 \text{ and } \hat{p}_l = \frac{1}{2n}\sum_{i=1}^{n} G_{il}.$$

The $10^5$ variants are split into 2,000 non-overlapping panels containing $m = 50$ markers each. In type 1 error simulations, phenotypes are re-simulated 100 times and 100 marker panels are tested for association with the simulated phenotype. Overall, 5 tests are performed for each of the 6,000 marker panels, which results in 30,000 replicates used for type 1 error estimation. Model I in Equation 15 and II in 16 are used to generate the traits with $\gamma = 0$.

In power simulations, for each given setting, we only simulate the 100,000 markers once and select a panel of $m = 50$ markers to be causal and tested for association with the traits. Model I is used to generate $k$ traits, where only a subset of the traits are associated with the selected marker panel. This is because it is unlikely in real applications for all the traits to be associated with the same set of markers. For $k = 2, 5, 50, 100$, we randomly choose 1, 2, 5 and 10 of the traits to be associated with the chosen marker panel, respectively. For each associated trait $j$, $\boldsymbol{\gamma}_j$ in Model I is chosen using the following conditions (independently across associated traits): (1) The overall effect of the $m$ markers explain 2.5% (if $k \leq 5$) or 3% (if $k > 5$) of the total trait variance; (2) Half of the $m$ markers are chosen at random to be causal, the rest are null; (3) Among the causal markers, half of the markers have positive effects and the other half have negative effects. For the remaining traits, which are not associated with the marker panel, $\gamma_j$ is set to 0. Phenotypes are re-simulated 1,000 times and the marker panel is tested for association with each phenotype replicate. This results in 1,000 replicates used for power estimation. For both type 1 error and power simulations, the two major genes and the ancestry proportion included in the Models I and II are assumed to be unobserved and so are not included as covariates when testing for association.

#### Supplemental Tables

**Table S1:** Genes with the Strongest Associations with Expression Levels across the 11 KEGG Pathways in Framingham Heart Study.

| | | | SINGLE | | | | LOGO | | | ALL | | |
|---|---|---|---|---|---|---|---|---|---|---|---|---|
| Pathway | Gene[a] | Chr | $T_2$-newperm | $T_2$-asymp | $T_1$-newperm | $T_1$-asymp | $T_2$-newperm | $T_1$-newperm | $T_1$-asymp | $T_2$-newperm | $T_1$-newperm | $T_1$-asymp |
| hsa04612 | **KLRC1** | 12 | 3.57e-43 | 1.57e-38 | 4.93e-43 | 1.00e-30 | 1.84e-31 | 6.84e-66 | 4.78e-60 | 4.74e-38 | 3.90e-73 | 5.49e-66 |
| | **KLRD1** | 12 | 3.21e-11 | 2.74e-10 | 1.73e-10 | 3.31e-09 | 1.07e-22 | 3.03e-43 | 1.50e-41 | 5.69e-37 | 8.28e-71 | 1.33e-64 |
| | KLRC3 | 12 | 1.49e-78 | 9.12e-70 | 7.99e-79 | 8.68e-56 | 8.80e-16 | 2.25e-28 | 2.17e-28 | 1.14e-36 | 7.43e-71 | 1.88e-64 |
| | HSP90AA1 | 14 | 3.13e-22 | 2.71e-20 | 7.85e-22 | 8.87e-18 | 5.06e-01 | 6.36e-01 | 3.91e-01 | 7.27e-14 | 4.87e-16 | 8.41e-17 |
| | **IFNG** | 12 | 1.13e-22 | 3.74e-19 | 3.24e-23 | 7.88e-17 | 4.01e-01 | 4.19e-01 | 2.12e-01 | 3.77e-07 | 1.06e-08 | 2.29e-09 |
| | **TNF** | 6 | 3.13e-06 | 6.41e-05 | 6.14e-06 | 7.81e-05 | 1.13e-04 | 1.22e-04 | 3.09e-03 | 1.36e-06 | 3.17e-06 | 2.33e-04 |
| | **HSPA1A** | 6 | 1.86e-01 | 2.57e-01 | 2.52e-01 | 2.97e-01 | 3.23e-06 | 5.45e-06 | 2.80e-04 | 4.53e-06 | 7.90e-06 | 3.86e-04 |
| | **HSPA1L** | 6 | 8.50e-03 | 2.15e-02 | 1.21e-02 | 2.40e-02 | 7.65e-05 | 9.96e-05 | 2.31e-03 | 4.67e-06 | 8.07e-06 | 3.93e-04 |
| | HSPA1B | 6 | 8.36e-03 | 2.28e-02 | 1.26e-02 | 2.45e-02 | 5.63e-05 | 7.51e-05 | 1.89e-03 | 8.30e-06 | 1.61e-05 | 6.40e-04 |
| | IFI30 | 19 | 8.42e-12 | 4.29e-11 | 8.29e-12 | 1.68e-10 | 1.03e-01 | 3.46e-01 | 6.72e-02 | 2.94e-05 | 2.47e-07 | 1.10e-09 |
| hsa03320 | **SCP2** | 1 | 9.78e-133 | 2.14e-111 | 1.80e-132 | 3.00e-95 | 9.99e-01 | 9.97e-01 | 9.78e-01 | 6.68e-26 | 6.42e-93 | 1.35e-74 |
| | **CPT2** | 1 | 5.40e-01 | 5.44e-01 | 5.19e-01 | 4.93e-01 | 1.89e-21 | 7.16e-82 | 1.87e-69 | 1.76e-21 | 1.17e-81 | 2.45e-69 |
| | ACSL5 | 10 | 1.20e-42 | 7.70e-38 | 1.14e-41 | 3.89e-31 | 2.67e-01 | 1.97e-01 | 1.40e-01 | 1.91e-20 | 1.98e-27 | 1.39e-25 |
| | **CYP27A1** | 2 | 2.18e-141 | 3.80e-128 | 3.79e-153 | 2.36e-131 | 6.93e-01 | 6.92e-01 | 8.38e-01 | 3.39e-16 | 5.66e-155 | 6.52e-116 |
| | FADS2 | 11 | 1.59e-112 | 1.80e-97 | 1.71e-103 | 8.40e-73 | 3.95e-01 | 4.36e-01 | 2.40e-01 | 1.79e-15 | 6.42e-49 | 7.81e-44 |
| | **ACSL6** | 5 | 3.98e-26 | 5.73e-21 | 1.71e-29 | 2.70e-21 | 1.68e-02 | 1.88e-02 | 2.22e-01 | 4.13e-09 | 2.96e-18 | 3.42e-11 |
| hsa04330 | **PSEN1** | 14 | 2.56e-21 | 3.26e-19 | 9.19e-22 | 2.79e-17 | 1.32e-01 | 1.15e-01 | 6.79e-02 | 6.03e-14 | 5.93e-16 | 1.01e-15 |
| | NUMB | 14 | 4.56e-03 | 6.28e-03 | 1.77e-03 | 2.32e-03 | 8.24e-10 | 6.20e-11 | 2.28e-10 | 4.65e-11 | 7.24e-13 | 3.96e-12 |
| | ADAM17 | 2 | 1.06e-27 | 6.74e-24 | 6.53e-28 | 2.56e-20 | 1.22e-02 | 2.50e-02 | 1.04e-02 | 4.14e-10 | 2.55e-17 | 1.16e-16 |
| | **NCSTN** | 1 | 2.74e-19 | 5.12e-18 | 2.52e-18 | 1.70e-15 | 7.21e-01 | 7.61e-01 | 4.90e-01 | 6.28e-09 | 1.39e-16 | 3.38e-18 |
| | MAML3 | 4 | 2.61e-23 | 1.74e-21 | 2.20e-23 | 1.70e-18 | 8.21e-01 | 7.74e-01 | 4.49e-01 | 2.43e-05 | 2.86e-05 | 1.03e-06 |

[a] Genes in bold are those that have been previously associated with a disease that is among (or associated to) the ones listed for the corresponding pathway in the KEGG database. MIM numbers of genes: KLRC1 [MIM 161555], KLRD1 [MIM 602894], KLRC3 [MIM 602892], HSP90AA1 [MIM 140571], IFNG [MIM 147570], TNF [MIM 191160], HSPA1A [MIM 603012], HSPA1L [MIM 140559], HSPA1B [MIM 603012], IFI30 [MIM 604664], SCP2 [MIM 184755], CPT2 [MIM 600650], ACSL5 [MIM 605677], CYP27A1 [MIM 606530], FADS2 [MIM 606149], ACSL6 [MIM 604443], PSEN1 [MIM 104311], NUMB [MIM 603728], ADAM17 [MIM 603639], NCSTN [MIM 605254], MAML3 [MIM 608991] .

**Table S2:** Genes with the Strongest Associations with Expression Levels across the 11 KEGG Pathways in Framingham Heart Study.

| | | | SINGLE | | | | LOGO | | | ALL | | |
|---|---|---|---|---|---|---|---|---|---|---|---|---|
| Pathway | Gene[a] | Chr | $T_2$-newperm | $T_2$-asymp | $T_1$-newperm | $T_1$-asymp | $T_2$-newperm | $T_1$-newperm | $T_1$-asymp | $T_2$-newperm | $T_1$-newperm | $T_1$-asymp |
| hsa04060 | **IL1R2** | 2 | 2.14e-09 | 7.84e-09 | 7.43e-10 | 6.26e-08 | 4.41e-20 | 3.31e-194 | 7.21e-136 | 2.99e-25 | 2.01e-212 | 3.62e-148 |
| | IL1RL2 | 2 | 8.62e-01 | 8.77e-01 | 7.95e-01 | 8.18e-01 | 6.74e-24 | 1.43e-196 | 3.61e-135 | 4.71e-24 | 5.04e-196 | 1.09e-134 |
| | **IL1R1** | 2 | 3.03e-04 | 7.90e-04 | 2.09e-04 | 4.47e-04 | 3.62e-23 | 3.51e-195 | 1.69e-134 | 6.95e-24 | 1.18e-195 | 8.48e-135 |
| | IL1RL1 | 2 | 1.15e-49 | 1.51e-42 | 8.79e-50 | 1.31e-37 | 1.17e-17 | 8.69e-162 | 3.68e-111 | 5.18e-23 | 8.50e-193 | 3.75e-132 |
| | **IL18R1** | 2 | 6.41e-21 | 1.94e-17 | 1.10e-19 | 1.63e-14 | 5.64e-19 | 1.62e-184 | 1.10e-126 | 6.05e-23 | 1.38e-192 | 4.68e-132 |
| | **IL18RAP** | 2 | 1.05e-234 | 4.24e-206 | 4.45e-240 | 5.24e-177 | 2.50e-11 | 1.27e-32 | 1.37e-21 | 7.34e-23 | 5.42e-192 | 1.29e-131 |
| | **CCR2** | 3 | 5.44e-12 | 1.23e-10 | 1.48e-12 | 2.67e-10 | 7.08e-14 | 3.28e-66 | 6.45e-51 | 6.63e-16 | 1.89e-71 | 7.52e-55 |
| | CCR1 | 3 | 1.45e-33 | 3.40e-29 | 1.07e-32 | 1.20e-23 | 3.18e-10 | 1.31e-46 | 8.48e-36 | 2.18e-14 | 3.06e-66 | 1.21e-50 |
| | XCR1 | 3 | 8.17e-01 | 8.29e-01 | 8.59e-01 | 8.52e-01 | 3.01e-13 | 8.08e-56 | 5.98e-42 | 4.76e-13 | 1.64e-55 | 1.04e-41 |
| | CCR3 | 3 | 9.72e-105 | 3.13e-93 | 1.53e-101 | 1.16e-77 | 1.78e-05 | 1.87e-12 | 2.60e-09 | 1.32e-12 | 1.06e-57 | 7.06e-44 |
| | CXCR6 | 3 | 2.85e-02 | 3.73e-02 | 2.98e-02 | 3.28e-02 | 3.39e-11 | 4.77e-49 | 5.66e-37 | 2.51e-11 | 2.39e-49 | 3.49e-37 |
| | CCR9 | 3 | 1.90e-01 | 2.15e-01 | 3.62e-01 | 3.84e-01 | 6.20e-10 | 9.49e-44 | 5.61e-33 | 4.55e-10 | 5.81e-44 | 4.14e-33 |
| | CXCL2 | 4 | 2.92e-01 | 2.92e-01 | 3.07e-01 | 2.89e-01 | 5.26e-10 | 5.62e-26 | 9.67e-25 | 5.04e-10 | 4.38e-26 | 7.77e-25 |
| | **LTA** | 6 | 3.67e-01 | 4.61e-01 | 2.23e-01 | 2.86e-01 | 2.08e-08 | 1.24e-08 | 2.70e-03 | 2.57e-08 | 1.10e-08 | 2.63e-03 |
| | CXCL3 | 4 | 8.13e-01 | 8.24e-01 | 7.44e-01 | 7.56e-01 | 1.85e-08 | 5.18e-24 | 9.77e-22 | 2.71e-08 | 9.55e-24 | 1.66e-21 |
| | LTB | 6 | 5.50e-01 | 6.11e-01 | 4.06e-01 | 4.31e-01 | 3.50e-08 | 2.26e-08 | 3.07e-03 | 4.54e-08 | 2.96e-08 | 3.52e-03 |
| | **TNF** | 6 | 3.13e-06 | 6.41e-05 | 6.14e-06 | 7.81e-05 | 5.12e-07 | 2.36e-07 | 9.11e-03 | 4.56e-08 | 2.93e-08 | 3.48e-03 |
| | IL7R | 5 | 8.62e-60 | 3.89e-54 | 9.63e-59 | 9.56e-46 | 4.26e-01 | 7.15e-01 | 4.19e-01 | 7.75e-08 | 4.57e-21 | 8.81e-21 |
| | **CXCL5** | 4 | 3.66e-22 | 1.14e-19 | 7.23e-22 | 6.18e-17 | 2.01e-03 | 2.03e-13 | 1.60e-11 | 3.57e-06 | 1.24e-18 | 8.50e-16 |
| | PPBP | 4 | 3.18e-01 | 3.28e-01 | 2.38e-01 | 2.08e-01 | 9.13e-06 | 1.05e-17 | 6.28e-15 | 5.49e-06 | 4.14e-18 | 2.86e-15 |
| | PF4 | 4 | 3.07e-01 | 3.18e-01 | 2.93e-01 | 2.87e-01 | 1.82e-05 | 9.04e-17 | 5.26e-14 | 1.39e-05 | 6.22e-17 | 4.01e-14 |
| | **IL22** | 12 | 1.82e-04 | 5.11e-04 | 2.12e-04 | 4.93e-04 | 3.86e-05 | 1.83e-08 | 2.45e-09 | 1.96e-05 | 1.20e-08 | 1.67e-09 |
| | PF4V1 | 4 | 2.88e-57 | 9.34e-51 | 5.43e-54 | 6.62e-42 | 1.68e-01 | 5.41e-02 | 7.45e-02 | 2.53e-05 | 3.10e-16 | 1.85e-13 |
| | **CXCL1** | 4 | 5.40e-06 | 2.98e-05 | 1.08e-05 | 5.38e-05 | 3.71e-04 | 3.90e-14 | 8.72e-12 | 2.74e-05 | 3.28e-16 | 1.96e-13 |
| | CXCL6 | 4 | 3.95e-01 | 4.21e-01 | 3.47e-01 | 3.64e-01 | 2.90e-05 | 5.73e-16 | 3.24e-13 | 3.30e-05 | 6.60e-16 | 3.75e-13 |
| | **IL26** | 12 | 1.58e-27 | 4.00e-23 | 3.57e-27 | 2.82e-19 | 9.67e-04 | 8.09e-06 | 7.76e-07 | 3.98e-05 | 3.15e-08 | 3.50e-09 |
| | TNFSF8 | 9 | 2.13e-10 | 1.71e-09 | 1.34e-10 | 1.88e-09 | 6.38e-04 | 6.30e-03 | 1.48e-03 | 4.03e-05 | 2.58e-04 | 5.52e-05 |

[a] Genes in bold are those that have been previously associated with a disease that is among (or associated to) the ones listed for the corresponding pathway in the KEGG database. MIM numbers of genes: IL1R2 [MIM 147811], IL1RL2 [MIM 604512], IL1R1 [MIM 147810], IL1RL1 [MIM 601203], IL18R1 [MIM 604494], IL18RAP [MIM 604509], CCR2 [MIM 601267], CCR1 [MIM 601159], XCR1 [MIM 600552], CCR3 [MIM 601268], CXCR6 [MIM 605163], CCR9 [MIM 604738], CXCL2 [MIM 139110], LTA [MIM 153440], CXCL3 [MIM 139111], LTB [MIM 600978], TNF [MIM 191160], IL7R [MIM 146661], CXCL5 [MIM 600324], PPBP [MIM 121010], PF4 [MIM 173460], IL22 [MIM 605330], PF4V1 [MIM 173461], CXCL1 [MIM 155730], CXCL6 [MIM 138965], IL26 [MIM 605679], TNFSF8 [MIM 603875] .

**Table S3:** Genes with the Strongest Associations with Expression Levels across the 11 KEGG Pathways in Framingham Heart Study.

| | | | SINGLE | | | | LOGO | | | ALL | | |
|---|---|---|---|---|---|---|---|---|---|---|---|---|
| Pathway | Gene[a] | Chr | $T_2$-newperm | $T_2$-asymp | $T_1$-newperm | $T_1$-asymp | $T_2$-newperm | $T_1$-newperm | $T_1$-asymp | $T_2$-newperm | $T_1$-newperm | $T_1$-asymp |
| hsa04630 | PIK3R3 | 1 | 9.87e-64 | 2.14e-52 | 4.89e-65 | 8.74e-48 | 6.01e-03 | 8.97e-03 | 1.33e-01 | 6.21e-19 | 1.20e-32 | 1.38e-21 |
| | IL7R | 5 | 8.62e-60 | 3.89e-54 | 9.63e-59 | 9.56e-46 | 1.50e-01 | 9.95e-02 | 3.95e-02 | 3.20e-17 | 1.47e-45 | 6.60e-43 |
| | **IFNGR2** | 21 | 4.61e-06 | 1.40e-05 | 3.18e-05 | 5.03e-05 | 1.04e-03 | 8.00e-04 | 3.38e-06 | 2.96e-05 | 3.07e-06 | 3.57e-09 |
| hsa05321 | **IL18RAP** | 2 | 1.05e-234 | 4.24e-206 | 4.45e-240 | 5.24e-177 | 1.70e-05 | 9.17e-07 | 1.72e-05 | 8.84e-35 | 9.78e-249 | 1.51e-182 |
| | **IL18R1** | 2 | 6.41e-21 | 1.94e-17 | 1.10e-19 | 1.63e-14 | 2.38e-22 | 3.60e-226 | 9.75e-167 | 1.52e-34 | 2.45e-248 | 2.54e-182 |
| | **NOD2** | 16 | 4.14e-163 | 3.72e-148 | 1.84e-152 | 1.10e-115 | 4.48e-01 | 5.28e-01 | 1.72e-01 | 1.16e-33 | 8.14e-99 | 1.16e-93 |
| | **TLR5** | 1 | 1.89e-24 | 1.46e-21 | 2.77e-24 | 9.01e-18 | 3.94e-01 | 6.93e-01 | 4.02e-01 | 9.47e-11 | 6.13e-07 | 3.60e-08 |
| | TLR4 | 9 | 2.93e-34 | 3.56e-30 | 1.49e-33 | 1.26e-24 | 1.95e-01 | 2.31e-01 | 1.30e-01 | 7.35e-10 | 8.19e-18 | 1.05e-17 |
| | **IL23R** | 1 | 1.37e-03 | 2.36e-03 | 1.31e-03 | 1.40e-03 | 6.26e-06 | 1.10e-05 | 9.19e-06 | 4.28e-07 | 3.34e-07 | 3.27e-07 |
| | **IL12RB2** | 1 | 1.59e-11 | 2.27e-10 | 4.56e-11 | 1.13e-09 | 7.13e-03 | 1.52e-02 | 7.41e-03 | 5.20e-06 | 6.75e-06 | 3.07e-06 |
| | IL22 | 12 | 1.82e-04 | 5.11e-04 | 2.12e-04 | 4.93e-04 | 3.82e-05 | 7.57e-07 | 1.11e-07 | 5.38e-06 | 1.37e-07 | 2.08e-08 |
| | **TNF** | 6 | 3.13e-06 | 6.41e-05 | 6.14e-06 | 7.81e-05 | 1.67e-03 | 1.84e-03 | 2.51e-02 | 1.66e-05 | 3.53e-05 | 1.67e-03 |
| hsa04140 | PIK3R3 | 1 | 9.87e-64 | 2.14e-52 | 4.89e-65 | 8.74e-48 | 1.60e-01 | 4.46e-02 | 2.29e-01 | 1.26e-19 | 5.83e-36 | 3.72e-25 |
| | DAPK1 | 9 | 6.31e-149 | 1.79e-126 | 1.82e-149 | 1.85e-98 | 2.63e-01 | 1.81e-01 | 8.03e-02 | 1.88e-15 | 1.52e-81 | 2.07e-71 |
| | VAMP8 | 2 | 3.07e-32 | 4.93e-27 | 2.31e-30 | 3.43e-22 | 6.87e-03 | 2.83e-03 | 3.28e-04 | 4.69e-12 | 4.49e-19 | 3.54e-19 |
| | MAPK3 | 16 | 2.21e-17 | 2.59e-16 | 2.74e-17 | 2.97e-15 | 9.06e-01 | 8.02e-01 | 4.62e-01 | 2.41e-10 | 5.71e-19 | 1.47e-20 |
| | CTSB | 8 | 1.30e-04 | 1.37e-03 | 3.78e-07 | 8.82e-05 | 1.67e-04 | 7.03e-05 | 6.61e-02 | 1.19e-05 | 1.93e-08 | 1.95e-03 |
| hsa04022 | MAPK3 | 16 | 2.21e-17 | 2.59e-16 | 2.74e-17 | 2.97e-15 | 4.82e-01 | 7.13e-01 | 3.21e-01 | 9.96e-10 | 2.91e-15 | 4.80e-17 |
| | TRPC6 | 11 | 1.57e-37 | 2.73e-31 | 9.81e-38 | 5.27e-26 | 7.57e-02 | 1.49e-01 | 2.98e-01 | 1.94e-07 | 5.76e-10 | 1.23e-07 |

[a] Genes in bold are those that have been previously associated with a disease that is among (or associated to) the ones listed for the corresponding pathway in the KEGG database. MIM numbers of genes: PIK3R3 [MIM 606076], IL7R [MIM 146661], IFNGR2 [MIM 147569], IL18RAP [MIM 604509], IL18R1 [MIM 604494], NOD2 [MIM 605956], TLR5 [MIM 603031], TLR4 [ MIM 603030], IL23R [MIM 607562], IL12RB2 [MIM 601642], IL22 [MIM 605330], TNF [MIM 191160], DAPK1 [MIM 600831], VAMP8 [MIM 603177], MAPK3 [MIM 601795], CTSB [MIM 116810], TRPC6 [MIM 603652] .
